# Reconstructing Spatio-Temporal Trajectories of Visual Object Memories in the Human Brain

**DOI:** 10.1101/2022.12.15.520591

**Authors:** Julia Lifanov, Benjamin J. Griffiths, Juan Linde-Domingo, Catarina S. Ferreira, Martin Wilson, Stephen D. Mayhew, Ian Charest, Maria Wimber

**Affiliations:** School of Psychology and Centre for Human Brain Health (CHBH), University of Birmingham, Birmingham, United Kingdom; Mind, Brain and Behavior Research Center, University of Granada, Granada, Spain; Department of Experimental Psychology, University of Granada, Granada, Spain; Research Group Adaptive Memory and Decision Making, Max Planck Institute for Human Development, Berlin, Germany; Center for Adaptive Rationality, Max Planck Institute for Human Development, Berlin, Germany; Max Planck Dahlem Campus of Cognition, Max Planck Institute for Human Development, Berlin, Germany; Institute of Health and Neurodevelopment (IHN), School of Psychology, Aston University, Birmingham, United Kingdom; Département de Psychologie, Université de Montréal, Montréal, Quebec, Canada; School of Psychology & Neuroscience and Centre for Cognitive Neuroimaging (CCNi), University of Glasgow, Glasgow, United Kingdom

**Author notes:** **Contact Information / Corresponding Authors** Maria Wimber, Julia Lifanov.

## Abstract

Our understanding of how information unfolds when we recall events from memory remains limited. In this study, we investigate whether the reconstruction of visual object memories follows a backward trajectory along the ventral visual stream with respect to perception, such that their neural feature representations are gradually reinstated from late areas close to the hippocampus backwards to lower-level sensory areas. We use multivariate analyses of fMRI activation patterns to map high-and low-level features of the object memories onto the brain during retrieval, and EEG-fMRI fusion to track the temporal evolution of the reactivated patterns. Participants studied new associations between verbs and randomly paired object images in an encoding phase, and subsequently recalled the objects when presented with the corresponding verb cue. Decoding reactivated memory features from fMRI activity revealed that retrieval patterns were dominated by conceptual features, represented in comparatively late visual and parietal areas. Representational-similarity-based fusion then allowed us to map the EEG patterns that emerged at each given time point of a trial onto the spatially resolved fMRI patterns. This fusion suggests that memory reconstruction proceeds from anterior fronto-temporal to posterior occipital and parietal regions, partly in line with a semantic-to-perceptual gradient. A linear regression on the peak time points of reactivated brain regions statistically confirms that the temporal progression is reversed with respect to encoding in ventral stream areas. Together, the results shed light onto the spatio-temporal trajectories along which memories and their constituent features are reconstructed during associative retrieval.

**Significance statement:** In our work, we combined EEG and fMRI to investigate which features of a visual object are reactivated when recalled from episodic memory, and how the memory reconstruction stream unfolds over time and across the brain. Our findings suggest that with respect to perception, memory retrieval follows a backwards information trajectory along a conceptual-to-perceptual gradient, and additionally relays retrieved information to multimodal fronto-parietal brain regions. These findings address the fundamental question of whether memories are more or less complete and truthful reconstructions of past events, or instead are subject to systematic biases that prioritise some types of features over others. Our data suggests that episodic memory retrieval is a dynamic and highly reconstructive process with clear prioritisation of abstract-conceptual over detailed-perceptual information.

## Introduction

Relative to the substantial literature on visual perception, little is known about how the human brain reconstructs content that is retrieved from memory. We here investigated which features are preferentially reactivated where in the brain and when in time when visual objects are recalled from episodic memory.

Visual object perception triggers an information processing cascade along the ventral visual stream which, in the first few hundred milliseconds, follows a gradient of increasing abstraction: Early processing stages are dominated by low-level perceptual representations, while later stages increasingly code for conceptual features (Carlson et al., 2013; Cichy et al., 2014; Clarke & Tyler, 2015; Kolb et al., 1995; Martin et al., 2018). After approximately 300ms, information reaches the hippocampus, where concept cells represent objects as highly abstract, perceptually invariant concepts (Quiroga, 2012). The hippocampus presumably binds together information from different sources (including dorsal stream) and serves the longer-term storage and retrieval of associative information (Danker et al., 2017; Eichenbaum, 2001; Rolls, 2010).

Computational models assume that the hippocampus can then use a partial reminder to recover the missing, associatively linked information, a process termed pattern completion (Marr, 1971; O’Reilly & McClelland, 1994; Rolls, 2010, 2013; Teyler & DiScenna, 1986). Pattern completion in turn dictates that neocortical brain regions re-instantiate the various features of the relevant event (Marr, 1971; Teyler & Rudy, 2007; Teyler & DiScenna, 1986; Rolls, 2013; also see Moscovitch, 2008), presumably leading to the subjective re-experiencing of a past event (Tulving et al., 1983).

Central to these computational models, they describe memory reinstatement as an all-or-none process and assume that during memory recall, neocortical information is re-established the way it was represented during initial perception. Indices for copy-like encoding-retrieval-similarities (ERS; Staresina et al., 2012) have been found by various neuroimaging studies using similarity-or classification-based multivariate analyses (see Danker & Anderson, 2010; Rissman & Wagner, 2012; Staresina et al., 2012), including the reinstatement of individual episodes, semantic category, sensory modality, or task context (Wing et al., 2014; Ferreira et al., 2019; Griffiths et al., 2019; Jiang et al., 2020; Polyn et al., 2005). Evidence for pattern reinstatement has been found throughout the sensory processing pathways activated during initial event perception, including areas proximal to the hippocampus (Staresina et al., 2012) as well as more distant neocortical structures (Polyn et al., 2005; Wing et al., 2014). Additionally, successful retrieval is associated with the interplay between hippocampal activity and cortical reinstatement (Bosch et al., 2014; Horner et al., 2015; Staresina et al., 2012). Together, these findings support the central premise that hippocampal pattern completion is associated with a more or less truthful reinstatement of encoding patterns. In the present study, our first goal is to decompose visual object memories into perceptual and conceptual components, and to test if both types of features are equally reinstated, in the same areas representing these features during encoding. The latter question is particularly interesting in the light of recent evidence that information is not necessarily decodable from the same regions during encoding and retrieval (Favila et al., 2020; Xue, 2022).

Moving beyond spatial localization we asked next how the neocortical reinstatement process unfolds over time, and whether different features of a memory are reactivated with different temporal priority. Such prioritisation could naturally emerge from the hippocampus’ differential connectivity with neocortical regions during retrieval. Once pattern completion is initiated, most information is relayed via the subiculum to the entorhinal cortex which in turn projects back to other cortical areas (Chrobak et al., 2000; Rolls, 2013; Staresina & Wimber, 2019). A majority of direct hippocampal and entorhinal efferents terminate in late areas of the visual processing hierarchy where representations cluster in abstract-semantic categories, while there are few connections to low-level sensory areas (Dalton et al., 2022; Insausti & Muñoz, 2001). We therefore expect that memories are initially reinstated on a conceptual level, and information coded in lower-level visual areas can only be reached at later stages of memory reconstruction (Insausti & Muñoz, 2001; Linde-Domingo et al., 2019). Indeed recent studies using EEG decoding and behavioural responses demonstrated a temporal prioritization of high-level conceptual over perceptual features during memory retrieval (Linde-Domingo et al., 2019; Lifanov et al., 2021), mirroring the hierarchy during perception of the same stimuli. Together with studies showing a similar reversed hierarchy and temporal prioritisation during mental imagery as opposed to perception (Dijkstra et al., 2020; Horikawa & Kamitani, 2017), these findings suggest that reconstructing an object from memory induces a feed-backward propagation along the ventral visual stream. Where in the brain different memory features are reinstated, and whether the observed conceptual-to-perceptual gradient of feature recall truly maps onto a sensory backpropagation stream, remains an open question that is central to our study.

The present study combined fMRI with EEG pattern analysis to test if recalling an object from memory elicits a backwards processing cascade relative to perception. An associative recall paradigm with visual object stimuli similar to previous studies (Linde-Domingo et al., 2019, Lifanov et al., 2021) allowed us to differentiate between low-level perceptual and high-level conceptual object features. First, using multivariate analysis of the fMRI patterns only, we compared where in the brain these low-and high-level feature dimensions can be decoded during retrieval as opposed to encoding. Second, we asked whether the object information reconstructed during retrieval propagates backwards along the ventral visual stream, from high-level semantic to low-level sensory areas, using EEG-fMRI fusion to resolve spatial activation patterns in time.

## Results

In our memory paradigm, based on Linde-Domingo et al. (2019), participants first encoded novel associations between objects and action verbs. Importantly, object images varied along a perceptual (photographs versus line drawings) and a conceptual (animate versus inanimate) dimension that were used for later classification to distinguish between the two hierarchical processing levels. At the recall stage, following a distracter task, participants were asked to retrieve the object in as much detail as possible when presented with the verb, and to indicate the time point of subjective recall with a button press while holding the retrieved image in mind. Following this subjective button press, participants answered either a perceptual (Was the object a photograph/drawing?) or conceptual (Was the object animate/inanimate?) question on each recall trial. Using this paradigm in an fMRI environment, we aimed at pinpointing the retrieval-related reinstatement of different object features in the brain. A searchlight classification approach was used to spatially locate perceptual (photograph versus drawing) and conceptual (animate versus inanimate) representations during image encoding and retrieval. The next aim was to map spatial representations onto a precise timeline preceding the subjective recall button press on a trial-by-trial basis within subjects, using simultaneous EEG-fMRI data acquisition. Since the within-scanner data turned out to be too noisy to decode the relevant object features (see ‘Analyses that did not work’), we instead used a previously acquired out-of-scanner EEG dataset (Linde-Domingo et al., 2019) to map the spatial representations onto the memory reconstruction timeline. This was done by means of an EEG-fMRI data fusion, using a second-order representational similarity analysis (RSA, Kriegeskorte, 2009; Kriegeskorte et al., 2006, 2008; Kriegeskorte & Kievit, 2013; Nili et al., 2014).

## Behaviour

We first analysed the behaviour of participants who completed the fMRI study and descriptively compared their performance with the participants who took part in the out-of-scanner EEG study (Linde-Domingo et al., 2019). The in-and out-of-scanner encoding data showed comparable RTs and accuracy rates (Table 1). However, looking at subjective retrieval RTs (Fig. 1d), participants in the fMRI study were 1.24 s faster on average to indicate subjective retrieval than participants in the EEG study. This RT difference was not only a result of the repeated retrievals in the fMRI group, since RTs of both the first (*M1* = 1.93 s; *SD1* = 0.67 s) and second (*M2* = 1.58 s, *SD2* = 0.66 s) retrieval repetition of the fMRI group were substantially shorter than those of the EEG group. Participants in the fMRI sample might thus have tended to indicate subjective retrieval prematurely, possibly due to discomfort in the scanner environment, and this may have had knock-on effect on the catch question RTs discussed in the next paragraph.

**Table 1.** Means (M) and Standard deviations (SD) of encoding and subjective retrieval reaction times (RTs) and accuracies of in-and out-of-scanner data.

|  | In-scanner | Out-of-scanner |
| --- | --- | --- |
| Encoding RT | $M = 3.03$ s; $SD = 0.90$ s | $M = 2.82$ s, $SD = 1.56$ s |
| Retrieval RT | $M = 1.75$ s; $SD = 0.66$ s | $M = 2.99$ s; $SD = .81$s |
| Retrieval accuracy | $M = 84.58$ %, $SD = .07$ | $M = 86.75$ %; $SD = .06$ |

**Figure 1.**
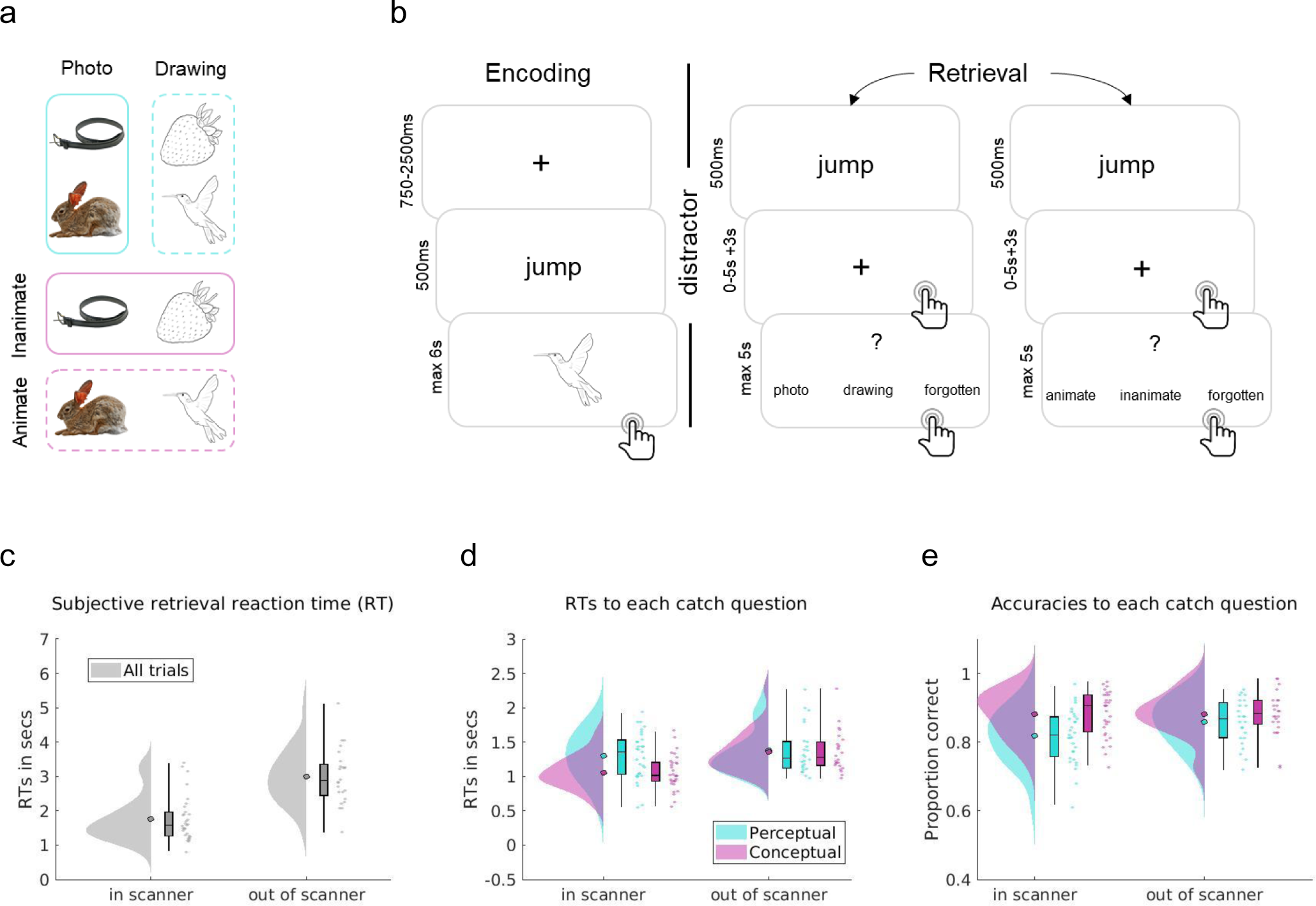
Overview of stimuli and task. a) Design of the stimuli. The 128 pictures used in any given participant were orthogonally split into 64 drawings and 64 photographs, out of which 32 were animate and 32 inanimate objects, respectively. Each object could thus be classified along a perceptual (photo/drawing, cyan) or conceptual (animate-inanimate, magenta) dimension. b) One prototypical task block of the paradigm. At encoding, participants were asked to associate verb-object pairings, and indicate the successful formation of an association by button press. After a 20 s distractor task, participants did a memory test where on each trial, they recalled the previously associated object upon presentation of a verb cue and indicated successful subjective recollection by button press (referred to as retrieval button press). Then participants were asked to hold the mental image of the object in mind for a further 3 seconds, before answering a perceptual or conceptual catch question. Each association was tested twice during retrieval, once with a perceptual, once with a conceptual question, with several intervening test trials in between repetitions of the same association. Participants performed 16 task blocks overall, with eight novel associations per block. Stimuli illustrated are chosen from the BOSS database (https://sites.google.com/site/bosstimuli/home, https://creativecommons.org/licenses/by-sa/3.0/) and customized with free and open source GNU image manipulation software (www.gimp.org; see Linde-Domingo et al., 2019). Figure adapted from Lifanov et al. (2021) and Linde-Domingo et al. (2019). c-e) Behavioural data acquired within (EEG-fMRI group) and out of scanner (EEG only group). c) Subjective retrieval reaction times (RTs), d) RTs for each type of catch question, and e) catch question accuracies for the two question types. Filled circles represent the overall mean, boxplots represent median and 25th and 75th percentiles; whiskers represent 2nd and 98th percentile; dots represent the means of individual subjects. Grey represents all trials, cyan represents responses to perceptual, magenta to conceptual questions. In-scanner data comes from n = 24 independent subjects, out-of-scanner-data from n = 31 independent subjects. Figures made using RainCloud plots Version 1.1 (https://github.com/RainCloudPlots/RainCloudPlots, Allen et al., 2019; Whitaker et al., 2019).

After an explorative inspection of the behavioural data, we compared accuracies and RTs between perceptual and conceptual catch questions using paired-sample t-tests (Table 2). Note that, different from our previous behavioural work (Linde-Domingo et al., 2019; Lifanov et al., 2021), the paradigm used here was optimised for capturing neural retrieval patterns rather than measuring feature-specific reaction times. After cue onset, participants were first instructed to mentally reinstate the object and indicate retrieval by button press. At this stage, we expected participants to have a fully reconstructed image of the recalled object in mind, such that answering the perceptual and conceptual catch question afterwards would take equally long. In the EEG study (also see Linde-Domingo et al., 2019), participants indeed answered conceptual and perceptual questions equally fast (*t*(23) = .5, *p* = .62 (uncorr.)) and accurately (*t*(23) = −1.80, *p* = .09 (uncorr.)). Participants in the fMRI study, however, performed less accurately (*t*(30) = −6.49, *p* < .01 (uncorr.)) and slower (*t* (30) = 8.47, *p* < .01 (uncorr.)) when answering perceptual compared to conceptual questions. The discrepancy between the EEG and fMRI samples, together with the difference in subjective retrieval time, suggests that participants in the fMRI study often pressed the subjective retrieval button before recall was complete, such that the feature reconstruction process carried over into the catch question period (Linde-Domingo et al., 2019; Lifanov et al., 2021).

**Table 2.** Means (M) and Standard deviations (SD) of reaction times (RTs) and accuracies for perceptual and conceptual questions separately of in-and out-of-scanner data and paired-sample t-test between perceptual and conceptual performances.

| a) In-scanner | RTs | Accuracy |
| --- | --- | --- |
| Perceptual | $M = 1.30$ s, $SD = 0.35$ s; | $M = 0.82$, $SD = 0.08$ |
| Conceptual | $M = 1.06$ s, $SD = 0.25$ s; | $M = 0.88$, $SD = 0.07$ |
| t-test | $t(30) = 8.47$,<br>$p < .01$ (uncorr.) | $t(30) = -6.49$,<br>$p < .01$ (uncorr.) |

| b) Out-of-scanner | RTs | Accuracy |
| --- | --- | --- |
| Perceptual | $M = 1.38$ s, $SD = 0.35$ s; | $M = 0.86$, $SD = 0.07$ |
| Conceptual | $M = 1.36$ s, $SD = 0.30$ s; | $M = 0.88$, $SD = 0.07$ |
| t-test | $t(23) = .5$,<br>$p = .62$ (uncorr.) | $t(23) = -1.80$,<br>$p = .09$ (uncorr.) |

Despite these RT differences, accuracies were comparably high in the two samples. Moreover, in the EEG-fMRI fusion analyses reported further below, the fMRI data is mainly used to derive the spatial representational patterns, while the precise time resolution is provided by the EEG data (for a review see Jorge et al., 2014). Considering the sluggishness of the hemodynamic response (Friston et al., 1994; Kruggel & von Cramon, 1999; Glover, 2011), the faster button presses in the fMRI sample should thus only have minimal effects on the fusion results.

## fMRI multivariate results

### fMRI searchlight LDA

Previous work suggests that during memory recall, conceptual information becomes accessible earlier than perceptual information, as evidenced in neural pattern decodability (Linde-Domingo et al., 2019) and behaviour (Lifanov et al., 2021; Linde-Domingo et al., 2019, and Fig. 1d). The first aim of the present study was to determine the locus of these perceptual and conceptual representations both during encoding and retrieval. We thus performed a searchlight-based, linear discriminant analysis (LDA). Two separate classifiers were trained to distinguish between the different perceptual (photographs versus drawings) and the different conceptual (animate versus inanimate) classes of our visual objects (see methods for details). During encoding (Fig. 2a), when the object was present on the screen, perceptual class could be decoded in posterior regions along the ventral visual stream including V1, V2, lingual and fusiform gyrus (*t*(30) = 4.98, *p* < .05 (FWE)). Conceptual class was decodable from more anterior lateral temporal lobe areas, including inferior, mid and superior temporal gyrus; and from precuneus, and inferior frontal gyrus, dorsolateral prefrontal cortex and anterior prefrontal cortex (*t*(30) = 4.82, *p* < .05 (FWE)). Together, this mapping generally reflects a posterior-to-anterior perceptual-to-conceptual processing gradient. The results largely mirror the univariate analyses reported in the supplemental materials (Fig. 2, Suppl. Fig. 1) and complement them by showing additional content decoding in frontal and parietal regions, suggesting that these areas code information in fine-grained multi-voxel patterns that univariate analyses are unable to capture.

**Figure 2.**
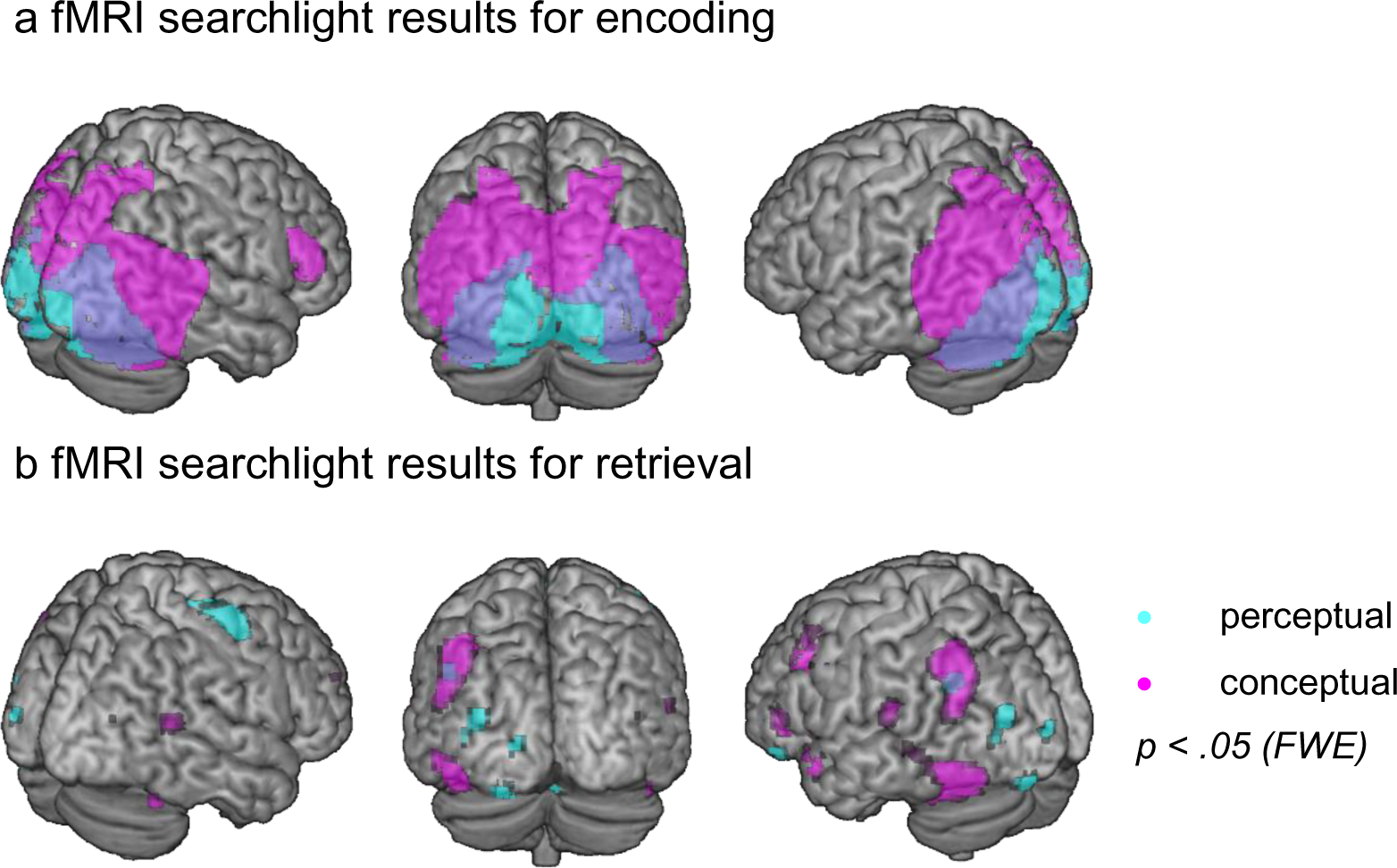
Searchlight LDA results. Second-level t-contrasts show a) Encoding and b) retrieval accuracies significantly higher than the 50% chance level when classifying perceptual (cyan) and conceptual (magenta) object features. All contrasts are thresholded at p < .05 (FWE-corrected). N = 31 independent subjects. Figure made using MRIcron (https://www.nitrc.org/projects/mricron, www.mricro.com, Rorden & Brett, 2000) and a colin 27 average brain template (http://www.bic.mni.mcgill.ca/ServicesAtlases/Colin27, Holmes et al., 1998; Copyright (C) 1993–2009 Louis Collins, McConnell Brain Imaging Centre, Montreal Neurological Institute, McGill University).

During retrieval (Fig. 2b), perceptual features were most strongly decodable in the precentral gyrus (i.e. premotor cortex), but also in precuneus, angular gyrus, cingulate cortex, inferior frontal gyrus and in ventral areas including V2, fusiform gyrus and middle temporal lobe (*t*(30) = 4.80, *p* < .05 (FWE)). Conceptual category membership was classified with the highest accuracy from fusiform gyrus, middle temporal gyrus, temporal pole, precuneus, angular gyrus, dorsolateral prefrontal cortex, and inferior and superior frontal gyrus (*t*(30) = 4.60, *p* < .05 (FWE); for detailed LDA results see tables 3-6 in the Supplementary file). Hence, in addition to the ventral visual areas dominating encoding/perception, we also found extensive frontal and parietal areas in our retrieval searchlight, in line with previous work suggesting that mnemonic as opposed to sensory content can often be decoded from these areas (Favila et al., 2018, 2020; Ferreira et al., 2019; Kuhl & Chun, 2014). Together, the fMRI-based classification of the two dimensions that we explicitly manipulated demonstrates that brain activity patterns during the retrieval of object images carry information about perceptual format and semantic content, with the latter dominating the recall patterns. Reinstatement was present in some of the areas seen at encoding, but additionally comprised regions in fronto-parietal networks.

No voxels survived (at p < .05, FWE-corrected) when cross-classifying perceptual or conceptual categories from encoding to retrieval, or from retrieval to encoding, which might be due to the overall noisier retrieval patterns or a transformation of these category representations between encoding and retrieval (see Discussion).

## EEG-fMRI data fusion

### EEG multivariate analysis

The next goal of this study was to map the spatial patterns in a given brain area onto the EEG patterns present at each time point of a trial, to reveal how content representations evolve over the time course of encoding and retrieval. For this data fusion, we first needed to obtain a meaningful EEG matrix representing object-to-object similarities for each time point of encoding and retrieval, which we could then compare to the fMRI-based matrices. Correlation-based RDMs for the EEG data were too noisy to see a meaningful match between modalities (see section ‘Analyses that did not work’). We thus created a classification-based matrix representing the dissimilarities between each pair of objects, in each EEG time bin, based on a cross-participant classifier. Amplitudes over the 128 electrodes at a given time point served as classification features, and our 24 participants provided the repetitions of each object. The pairwise classification accuracies were entered in a single, time resolved RDM structure representing the dissimilarity between individual objects of our stimulus pool collapsed across participants (Fig. 3a). For all fusion analyses, we investigated encoding timelines from image onset, and retrieval timelines leading up to the subjective button press.

**Figure 3.**
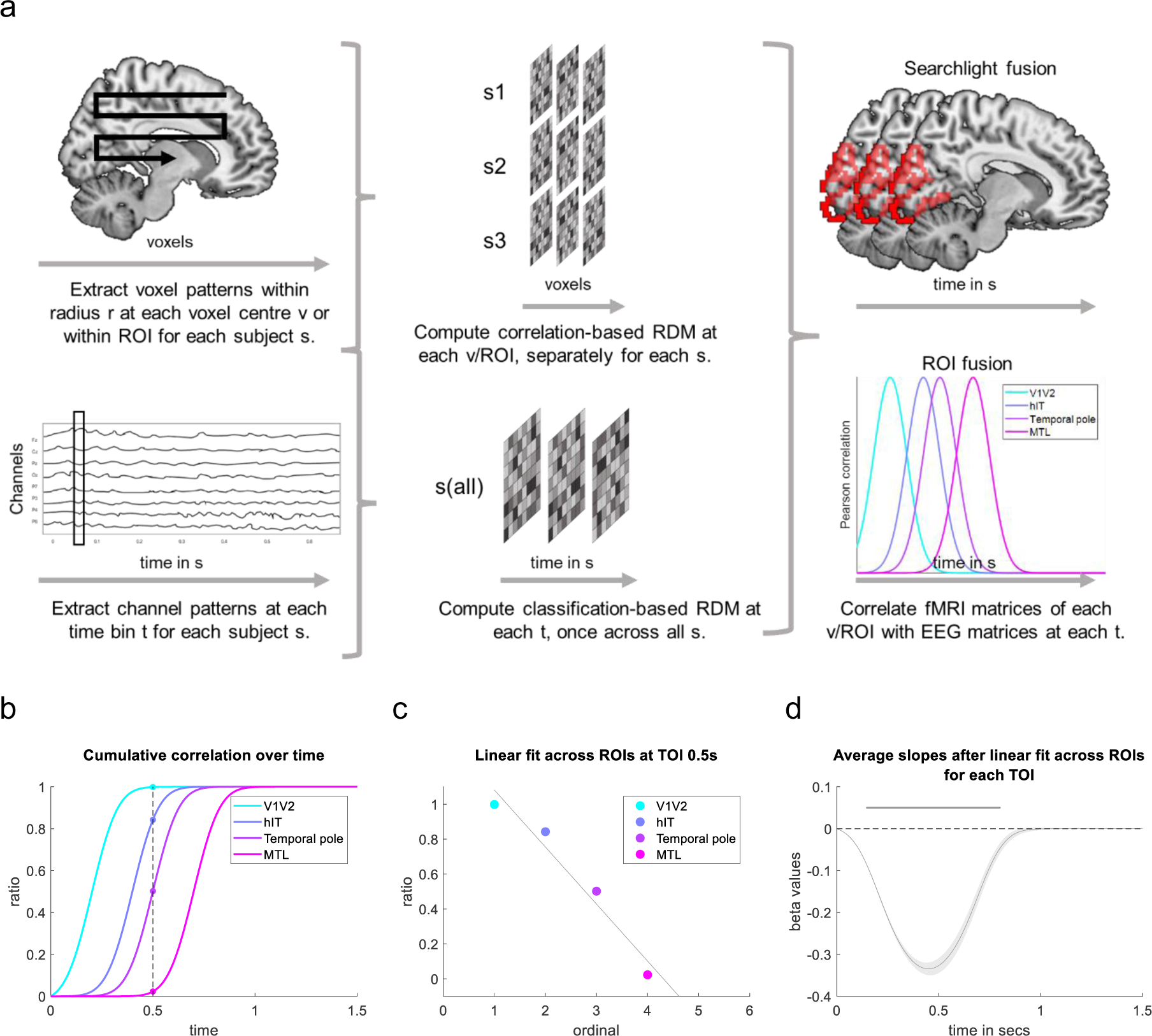
a) Overview of ROI and searchlight fusion. First, brain activity patterns are extracted from fMRI and EEG to create representational dissimilarity matrices (RDMs) in the spatial and temporal domain, respectively. Then, second-order correlations of the RDMs from the two imaging modalities are computed for the data fusion. The EEG RDMs are classification-based, derived from binary classifiers trained and tested to distinguish each pair of objects across participants. FMRI RDMs are computed within participants and are correlation-based. FMRI RDMs can be computed at each voxel centre (v) including voxels within a specific radius, or for larger regions of interest (ROI). Depending on this choice, the data fusion will be searchlight-or ROI-based, respectively. b-c) Linear regression approach on ROI time courses in a forward stream simulation. b) Normalized cumulative sum of each ROI time course of the ventral visual stream. At time point 0.5 s, earlier visual regions show a higher cumulative sum than later regions along the ventral visual stream. c) A linear regression is fitted across ROI values at 0.5 s and results in a negative slope. d) Average slopes and standard errors are illustrated for each time point across the window 0-1.5 s and tested against zero. Significant time points are indicated by points (p < .01 uncorrected). Method and panels b-d adapted from Michelmann et al. (2019). Brain figures made using MRIcron (https://www.nitrc.org/projects/mricron, www.mricro.com, Rorden & Brett, 2000), a colin 27 average brain template (http://www.bic.mni.mcgill.ca/ServicesAtlases/Colin27, Holmes et al., 1998; Copyright (C) 1993–2009 Louis Collins, McConnell Brain Imaging Centre, Montreal Neurological Institute, McGill University) and WFU PickAtlas v3.0 (Lancaster et al., 1997, 2000; Maldjian et al., 2003, 2004; Tzourio-Mazoyer et al., 2002).

### ROI fusion

Having obtained this EEG-based similarity time course, we performed a similarity-based fusion of the EEG and fMRI data, probing what brain regions (in the fMRI data) most closely matched the similarity structures contained in the EEG data at any given time point during encoding and retrieval. For a first fusion step, we used the representational geometries (i.e., the RDMs) from an anatomically pre-defined set of regions of interest (ROIs). We then calculated the correlation between these ROIs’ representational geometries and the geometries obtained from the EEG data for each time bin (Fig. 3a), and statistically compared the correlations against zero using cluster-basted permutation testing.

To statistically evaluate whether the information flow is primarily forward or backward along ventral visual stream areas, we used a linear regression on the cumulative sums of the correlation time courses of the corresponding ROIs, a method previously established for MEG analyses (Michelmann et al., 2019). The intuition of this method is that if region A starts accumulating information before region B, and region B before region C, the cumulative sums will line up sequentially to reflect this systematic delay in information accumulation (see simulation in Fig. 3b-d). In our case, we expected forward progressing latencies along the ventral visual stream during encoding and backward progressing latencies during retrieval (Fig. 3b-d).

Our predefined set of ROIs included regions along the ventral visual stream (V1/V2, hIT, temporal pole, and MTL), and regions along the dorsal stream (superior parietal lobe, lateral inferior parietal cortex, and medial parietal cortex; see section ROIs). Since many computational models propose that the role of the hippocampus is related to association or indexing rather than feature representation (Eichenbaum, 2001; McClelland et al., 1995; O’Reilly & Norman, 2002; Rolls, 2010, 2013; Teyler & DiScenna, 1986), we excluded the hippocampus from the MTL regions. In Fig. 4, correlation time courses are plotted for each individual ROI as t-values resulting from t-tests against zero, with clusters resulting from a cluster-based permutation test (see section ‘Analyses’). The figure shows ventral visual regions in panel a, and dorsal regions in panel b.

**Figure 4 a-d).**
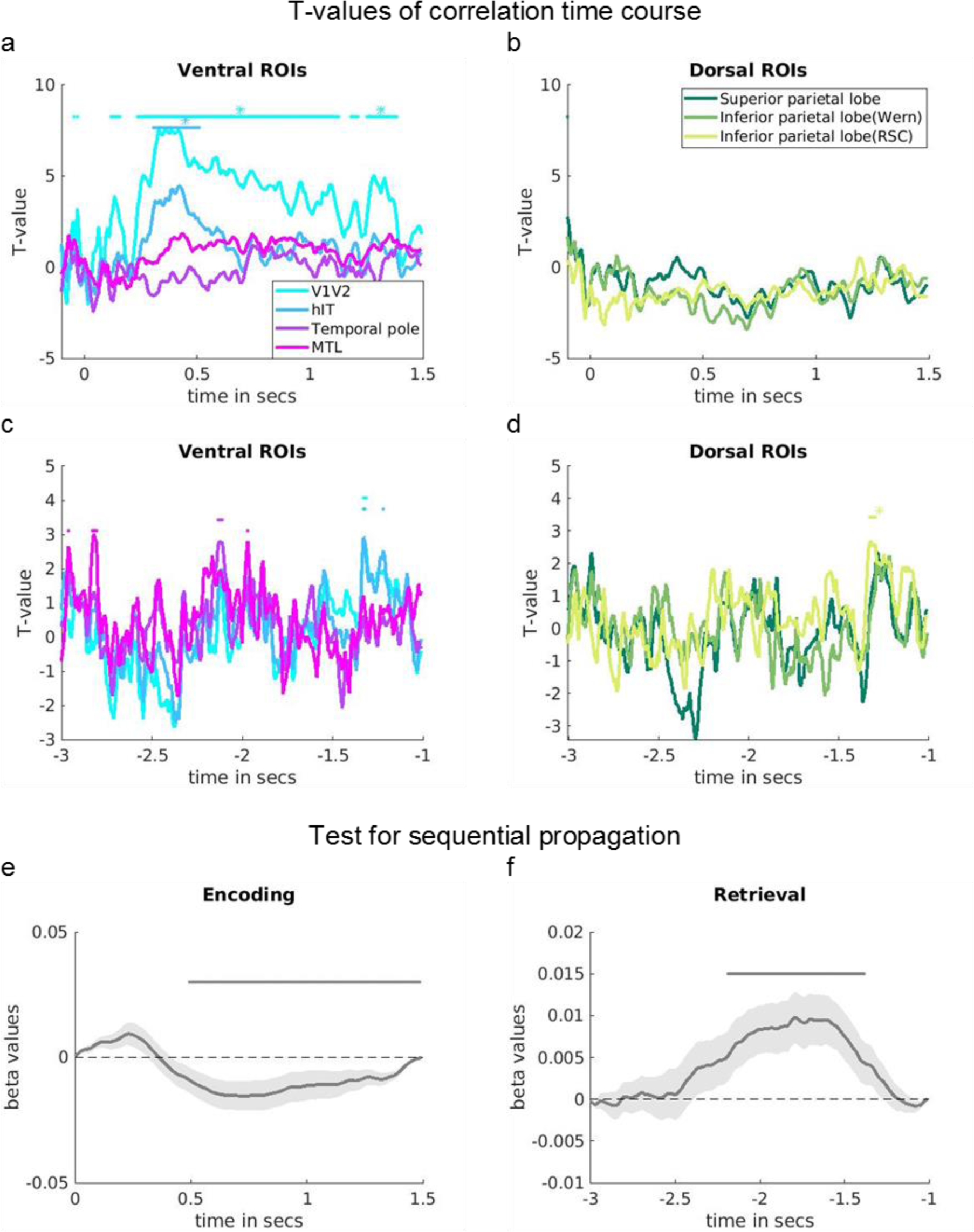
T-values from one-sample t-test of correlation time courses between the EEG RDMs and ventral visual (a & c) and dorsal (b & d) regions of interest (ROIs) during encoding (a & b) and retrieval (c & d). ROIs are colour-coded as indicated in the legends. Significant time points are indicated by points (p < .01 uncorrected) and asterisk (p < .05 cluster). e-f) Results of the test for sequential processing, using a linear fit of the cumulative sums of correlation time courses across ROIs within the ventral visual stream at e) encoding and f) retrieval. Average slopes and standard errors are shown for each time point and statistically tested against zero. Slopes are colour-coded as explained in the legends. A negative slope suggests that earlier ROIs along the ventral visual stream have a higher cumulative sum than later ROIs, indicative of a forward stream. According to the same logic, a positive slope indicates a backward stream. Significant time points are indicated by grey points above the curve (p < .05 cluster). Method adapted from Michelmann et al., 2019. In all subpanels, time point 0 s marks the object onset during encoding, and the subjective recall button press during retrieval. The latter is not included on time axis as it does not lie within the time window of interest (see Methods, also for details on cluster correction). Variance in all subpanels comes from n = 31 independent subjects. Brain figures made using MRIcron (https://www.nitrc.org/projects/mricron, www.mricro.com, Rorden & Brett, 2000), a colin 27 average brain template (http://www.bic.mni.mcgill.ca/ServicesAtlases/Colin27, Holmes et al., 1998; Copyright (C) 1993–2009 Louis Collins, McConnell Brain Imaging Centre, Montreal Neurological Institute, McGill University) and WFU PickAtlas v3.0 (Lancaster et al., 1997, 2000; Maldjian et al., 2003, 2004; Tzourio-Mazoyer et al., 2002).

Following the object onset at encoding (Fig. 4a-b), posterior regions including V1 and V2 showed an increasing correlation with the EEG representations from approximately 120 ms (p < .01 (uncorr.)). A significant cluster was seen from 240 ms onwards and another one at 1.31 s (p < .05 (cluster)). The first cluster was followed by a correlation increase of later ventral visual areas (human inferior temporal cortex (hIT)) around 310 ms (p < .01 (uncorr.)), reaching a significant peak at about 420 ms (p < .05 (cluster)). Later lateral and medial temporal as well as parietal regions did not show a significant correlation with the EEG time series during encoding.

As a proof of principle, we then tested for a forward processing stream in the encoding data across ventral visual areas. Performing a linear regression on the cumulative sums of all ventral ROI time courses at each time point (see Methods and Fig. 3b-c), we found negative slopes reaching significance 500 ms after stimulus onset (p < .05 (cluster)). This finding statistically corroborates the observation that earlier ventral visual regions code relevant information before later regions, supporting the well-established forward stream (Fig. 4e). It should be noted, however, that evidence for such feed-forward processing was only present relatively late in the trial, likely since the strongest peaks of object identity decoding also fell in this late time window (see Fig. 4, Suppl. Fig. 1).

When performing the same analysis for ventral visual ROIs during retrieval (Fig. 4c), several regions peaked below the threshold of cluster significance: MTL regions showed a peak correlation with the EEG representational geometries −2.96 s, −2.82 s and −1.97 s prior to the retrieval button press (p < .01 (uncorr.)). Further, correlation peaks were found for the temporal pole at −2.13 s (p < .01 (uncorr.)). HIT showed a peak correlation with the EEG geometry at −1.33 s and later at −1.22s (p < .01 (uncorr.)), with a peak by V1/V2 at −1.33 s before button press (p < .01 (uncorr.)).

Within the dorsal set of ROIs (Fig. 4d), the inferior medial parietal lobe showed a correlation peak in a similar time window, at −1.33 s before button press (p < .05 (cluster)). This peak in the inferior parietal, but no other parietal regions, coincided with the peak times of posterior ventral visual areas (V1/V2/hIT).

Our main aim of the data fusion was to test if memory reactivation followed a backward processing hierarchy along the ventral visual stream. We therefore also performed a linear regression on cumulative sums of ventral visual regions within retrieval. This analysis revealed a significant positive slope from −2.2 to −1.4 s before button press (p < .05 (cluster)), speaking in favour of a backward stream and thus confirming our main hypothesis (Fig. 4f).

### Searchlight fusion

Next, in a more explorative but also spatially better resolved approach, we conducted a whole-brain searchlight fusion to inspect where across the brain the fMRI-based representational geometries matched the EEG geometries over time. Figure 5 depicts spatial t-maps representing significant EEG-fMRI correlations in each searchlight radius for a given time bin, separately at encoding (Fig. 5a) and retrieval (Fig. 5b). The maps only show time points where significant spatial clusters emerged. During encoding, the searchlight fusion revealed a significant cluster in V1 and lingual gyrus at 60 ms following object onset, and again from 130 to 160 ms, and from 240 ms onwards, reactivating several more times (e.g. see timepoints 940 ms and 1.3 s) before fading around 1.4 s after stimulus onset. Importantly, over the time course of object perception, significant clusters of EEG-fMRI similarity gradually extended from early visual (V1, V2) to more ventral and lateral areas, including fusiform and inferior temporal gyrus. For the encoding data, the results of the searchlight approach thus overlap with the ROI fusion results, and mainly reveal ventral visual stream activation progressing in a forward (early-to-late visual cortex) manner.

**Figure 5.**
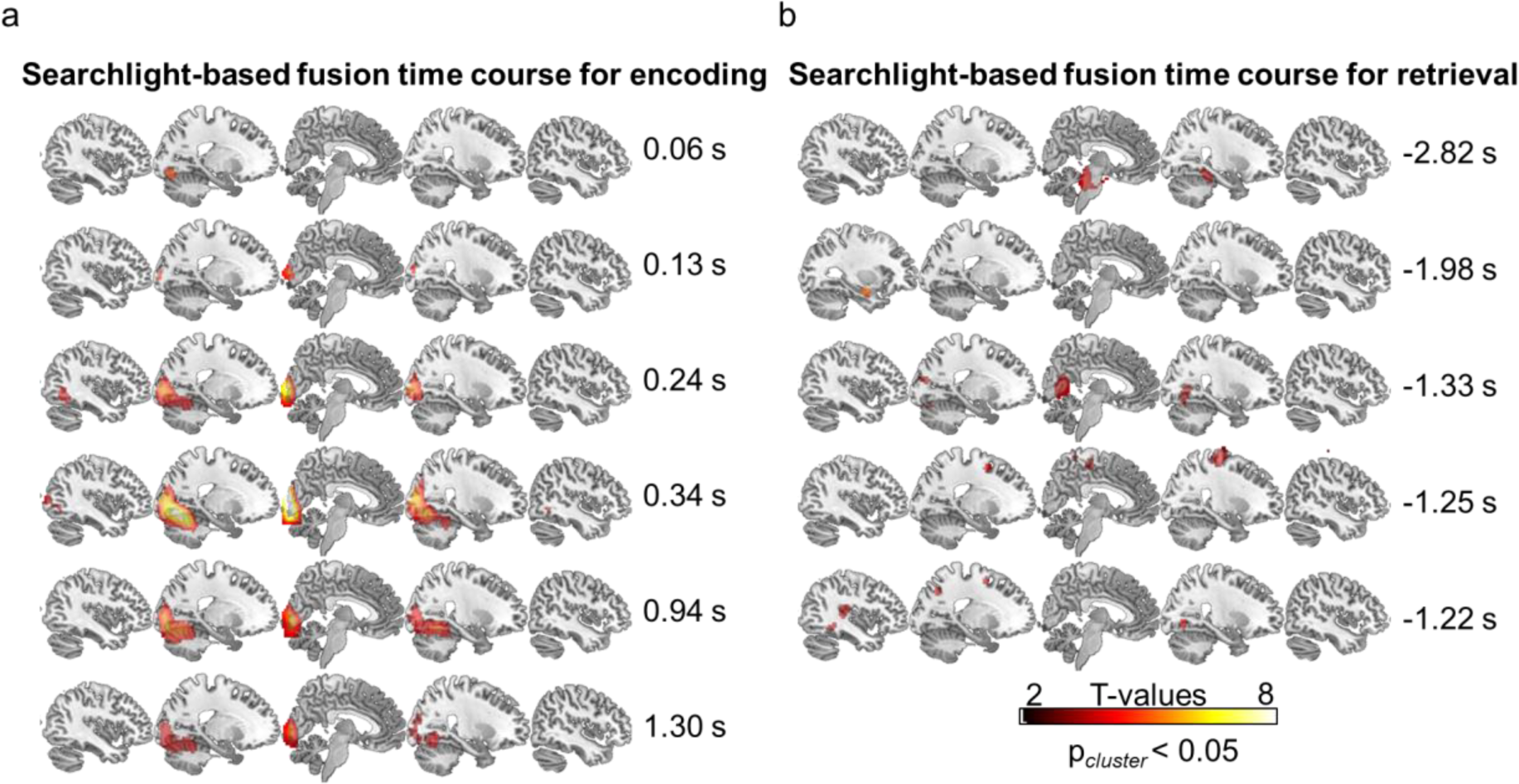
T-maps from one-sample t-test of EEG-fMRI correlations at a) encoding and b) retrieval showing time points in the trial time course where significant spatial clusters of correlations emerged (p_cluster_ < .05, see Methods for further details). T-test involved a spatial maximal permuted statistic correction combined with a threshold free cluster enhancement (Nichols & Holmes, 2002; Smith & Nichols, 2009). At encoding, time point 0 s would mark the object onset. At retrieval, time point 0 s marks the button press, however, both zero time points are not included in figure as no clusters emerged. N = 31 independent subjects. Brain figures made using MRIcron (https://www.nitrc.org/projects/mricron, www.mricro.com, Rorden & Brett, 2000) and a colin 27 average brain template (http://www.bic.mni.mcgill.ca/ServicesAtlases/Colin27, Holmes et al., 1998; Copyright Table 3. fMRI searchlight results for encoding: perceptual features.

During retrieval, the earliest cluster of significant EEG-fMRI correlations was found in the fusiform area, preceding the subjective retrieval button press by −2.82 s, accompanied by a small cluster in the pons. Contrary to our prediction, we also observed pattern overlap in hippocampus at −1.98 s before button press.

At time points closer to the button press, additionally clusters showing significant pattern correlations were found in lower-level regions. At −1.33 s, V1, V2, V4d and fusiform gyrus showed significant clusters. Further, at −1.25 s, clusters were found in premotor cortex, primary motor cortex, superior parietal lobule (SPL). Finally, clusters were found in premotor cortex, SPL, primary and associative visual cortex, fusiform area and Wernicke’s area at −1.22 s before subjective retrieval.

Together, the retrieval data showed convergence between the ROI-and searchlight fusion approach. It showed a progression of the peak correlations from fronto-temporal to posterior sensory regions, generally in line with a backward propagation hierarchy. This pattern of results is in line with previous work showing a conceptual-to-perceptual gradient in the same EEG dataset (Linde-Domingo et al., 2019). The present results additionally suggest that closest to the time point of subjective recollection (approximately −1.3 to −1.22 s preceding the button press), content information is most prominently represented in fronto-parietal and low-level visual areas.

## Discussion

What mnemonic content is recovered where in the brain during memory retrieval, and how does the hippocampal-neocortical pattern completion process unfold in time? Recent memory research suggests that the information processing hierarchy is reversed during the recall of a visual object from episodic memory compared with its initial perception (Linde-Domingo et al., 2019; Lifanov et al., 2021; Mirjalili et al., 2021), with conceptual features becoming available earlier than perceptual features. Here, we investigated the locus of these feature representations during encoding and recall using fMRI-based decoding. Additionally, EEG-fMRI fusion allowed us to test whether this presumed reversed information processing cascade during memory reconstruction maps onto the same ventral visual stream areas that carry the information forward during perception, but now following a backward trajectory. Our study shows that recall is largely dominated by conceptual information in the posterior ventral regions, and that the reinstatement of visual object memories follows a primarily backward trajectory along ventral visual object processing pathways, involving an additional information relay to parietal and frontal regions.

We first used uni-and multivariate analyses on the fMRI data alone to map out the regions processing perceptual and conceptual object dimensions that were built into the stimulus set. As expected, during encoding when the object was visible on the screen, these features largely mapped onto the ventral visual pathway, where early visual areas coded the perceptual features (coloured photos versus black-and-white line drawings), whereas later visual areas coded the mid-to higher-level conceptual information (animate versus inanimate objects; Long et al., 2018). This general perceptual-to-conceptual gradient is in line with a multitude of findings in the basic object recognition and vision literature (Carlson et al., 2013; Cichy et al., 2014; Kravitz et al., 2013).

Univariate analyses were not powerful enough to detect differential activations between perceptual and conceptual categories when objects were reconstructed from memory. Activity patterns at the finer-grained multi-voxel level, however, carried information about the categorical features of the retrieved images. Memory-related reactivation of these features comprised some regions also found during object encoding, particularly in late visual areas. This overlap confirms a body of previous work showing that posterior visual areas are not only involved in visual perception but also in internally generated processes, such as mental imagery (Dijkstra et al., 2021; Kosslyn et al., 1993) or episodic memory. While most memory studies find reactivation in late areas along the ventral stream (e.g., Griffiths et al., 2019; Polyn et al., 2005; Staresina et al., 2012), others have reported areas as early as V1 (Bosch et al., 2014). It is also commonly observed that stronger reinstatement is associated with memory success and strength (e.g. Huijbers et al., 2011; Ritchey et al., 2013; Staresina et al., 2012; Thakral et al., 2015; Wing et al., 2014) and the vividness and detail of remembering (Bone et al., 2020; Simons et al., 2022; St-Laurent et al., 2015; Wheeler et al., 2000), suggesting an important functional role of sensory reactivation. Assuming a hierarchical reverse processing from higher to lower sensory regions during recall, incomplete retrieval would lead primarily to a loss of perceptual information. Premature retrieval button presses, as found in the present study (see behavioural results), could indicate such incomplete recall and explain the overall weaker decodability of perceptual features.

Although the spatial localizations of feature-specific patterns during recall partly overlapped with those found during encoding, they also encompassed regions outside the ventral visual object processing pathways. Most notably, conceptual object information during recall was decodable from frontal and parietal regions. The role of fronto-parietal cortex in representing mnemonic information is still debated (Buckner et al., 1999; Favila et al., 2018, 2020; Humphreys et al., 2021; Kramer et al., 2005; Levy, 2012; Staresina & Wimber, 2019; Xue, 2022). A shift of content decoding from sensory areas during perception to multimodal fronto-parietal areas during memory recall is a frequent observation, and has recently been discussed as indicating a representational transformation (Favila et al., 2018, 2020; Xiao et al., 2017; Xue, 2022), potentially going along with a semanticization of the retrieved content (Ferreira et al., 2019; Lifanov et al., 2021). Some studies suggest that depending on the task and functional state (i.e. encoding or retrieval), content representations are preferentially expressed in sensory or parietal cortices, respectively, and mediated by differential connectivity with the hippocampus (Favila et al., 2018; Long & Kuhl, 2021; for a related review about the posterior medial network see Ritchey & Cooper, 2020). Others propose that parietal areas provide a contextualization of the retrieved memories (Jonker et al., 2018). Our findings support the previously described encoding-retrieval distinction and suggest that it is primarily conceptual information (Fig. 2b) that is represented in parietal regions during retrieval, in line with previous work showing strong object identity - and even abstract concept coding in parietal cortex (Ferreira et al., 2019; Jeong & Xu, 2016; Kaiser et al., 2022). The involvement of parietal networks is also discussed further below in relation to the timing of the reactivation.

Going beyond a purely spatial mapping of perceptual and conceptual representations, fusing the EEG and fMRI data allowed us to inject time information into these spatial maps, and to ask how the memory reconstruction stream evolves, building up to the time of subjective recollection. We used two complementary approaches for data fusion, both comparing the representational geometries found in the EEG patterns at each time point with the fMRI geometries found in a given brain region. Both the ROI-based and searchlight-based approach showed a mainly feedforward sweep of information processing during the first few hundred milliseconds of object perception, starting within early visual regions approximately 60 (searchlight fusion) to 120 ms (ROI fusion) after image onset, and spreading to more ventral visual regions within the next 200 ms. Importantly, a formal regression analysis allowed us to statistically corroborate the spatio-temporal direction of information flow, by modelling the information accumulation rate across the hierarchy of ventral visual stream regions (Michelmann et al., 2019). This regression analysis also showed a clear forward, posterior-visual to anterior-temporal progression of the correlation patterns during early phases of the encoding trials.

The EEG-fMRI fusion and sequence analysis applied to the retrieval data revealed a largely backward information processing trajectory, with information flowing from medial temporal lobe (both fusion approaches) and temporal pole (ROI fusion) to more posterior visual regions (both fusion approaches). The temporal pole has previously been associated with processing of high-level semantic information (e.g. Clarke, 2020; Noppeney & Price, 2002; Patterson et al., 2007; Rice et al., 2018; Visser et al., 2010). Moreover, temporal pole reactivation around −2.1 s before button press was close to the time point when objects from different conceptual classes were more discriminable than objects from similar conceptual classes based on the EEG classification alone (see Fig. 4, Suppl. Fig. 1b). Note, that our decoding peaks were found considerably earlier than in previous work (Linde-Domingo et al., 2019), likely due to the cross-participant object-identity decoding approach (for comparison see Linde-Domingo et al., 2019; Mirjalili et al., 2021) which may rely on higher-level exemplar-level information presumably processed close to the hippocampus (e.g. Clarke & Tyler, 2015). Together with our previous work using feature-specific reaction times (Linde-Domingo et al., 2019; Lifanov et al., 2021), these findings indicate that conceptual object features in semantic networks are among the first to be reactivated during memory retrieval.

More posterior areas including inferior temporal, inferior parietal and visual cortices, as well as frontal regions, reached their representational correlation peaks considerably later, from −1.33 s before button press onwards (Fig. 4c-d & 5b). While visual activations can be assumed to represent the perceptual content of the reactivated memories, the role of fronto-/parietal regions, as mentioned earlier, is still a matter of debate (e.g. Barry & Maguire, 2019; Fischer et al., 2021; Naghavi & Nyberg, 2005). In addition to the literature discussed above, parietal areas have been shown to play a role in contextual processing, imagery during recall and scene construction (Lundstrom et al., 2005; Fletcher et al., 1995; Chrastil, 2018). The latest reactivations might thus be indicative of a final stage of reinstatement leading up to subjective recollection, likely involving working memory and memory-related imagery, and hence, preparation for the upcoming categorisation task (Christophel et al., 2012, 2017; Ganis et al., 2004). In summary, contrary to encoding, we find that retrieval follows a backward propagating stream indicative of a conceptual-to-perceptual reconstruction gradient. Moreover, the backward reconstruction flow is not limited to ventral visual brain areas that dominate the encoding patterns, but instead involves frontal and parietal regions that may serve as an episodic memory or imagery buffer for the retrieved representations (Baddeley, 1998, 2000; Levy, 2012; Wagner et al., 2005).

The anterior-to-posterior retrieval stream shown here is reminiscent of a similar reversed stream for object imagery in absence of an episodic cue (Dijkstra et al., 2020), which has also been described as progressing in a hierarchical fashion along the ventral stream (Horikawa & Kamitani, 2017). Generating mental images from general knowledge compared to cuing them from episodic memory likely involve different neural sources, in the latter case most notably the hippocampus. However, both processes rely on the internal generation of stored mental representations (i.e., memories), and it is thus perhaps not surprising that mental imagery and episodic retrieval overlap in many aspects, including their reverse reconstruction gradient. While there have been various studies comparing imagery with perception (Dijkstra et al., 2020; Horikawa & Kamitani, 2017; Xie et al., 2020), it will be interesting for future studies to provide a more detailed picture of where in the processing hierarchy imagery and memory retrieval overlap.

The fMRI classification and the RSA-based EEG-fMRI fusion in this study offer complementary information. RSA is a highly useful tool to enable the comparison of neural representations measured by different neuroimaging modalities (Kriegeskorte & Kievit, 2013). By creating similarity structures from EEG and fMRI activity patterns in the same format, it becomes possible to directly correlate the representational geometries emerging in neural space and time. However, RSA is in itself blind to the informational content that is driving the match in representations (see Schyns et al., 2020). Classification-based approaches on fMRI (or EEG) data, on the other hand, can pinpoint categorical representations that are explicitly built into the experimental design. However, they are restricted to very few categories (e.g., animacy, colour), in turn limiting the conclusions that can be drawn about the certainly much richer and more multidimensional content of mental (memory) representations. In sum, RSA and categorical decoding methods offer complementary insights into the content of reactivated memories, likely explaining why in our study, some correlation peaks in the searchlight fusion maps during retrieval did not overlap with the sources of either perceptual or conceptual features. Richer study designs with more stimulus variations on a trial-by-trial basis (Schyns et al., 2020) are a promising avenue for revealing what type of content representations are contained in, and dominate, reactivated memories. An exciting recent development is the use of deep neural networks (DNNs) trained on image categorization, and to compare the networks’ layer patterns to the representational patterns that emerge in the real brain (Allen et al., 2022; Cichy et al., 2019). Some recent work has employed this approach to shed light onto the nature of reactivated memory representations, for example to investigate what level of feature reactivation predicts the vividness and distinctiveness of a memory (Bone et al., 2020). Others have used DNNs in comparison with intracranial EEG to reveal memory transformations over time and across different processing stages (Liu et al., 2020). Finally, different types of models, including word embedding models, have been used to uncover the temporal and content structure of recall narratives, and how it compares to the original encoding (Heusser et al., 2021). In contrast to the relative black box of DNN layers, the latter models offer more transparency and interpretability of the dimensions contained in a memory, and a route to understanding how the recall of naturalistic, continuous events unfolds in time.

In summary, our study supports the idea of a conceptual-to-perceptual information processing gradient that is reversed with respect to encoding along ventral visual stream regions processing object identity. Additionally, we show that the backward trajectory of memory reconstruction is not limited to ventral visual stream regions involved during encoding, engaging multisensory fronto-parietal areas at distinct stages of retrieval processing. The present findings demonstrate how the fusion of temporally and spatially resolved methods can further our understanding of memory retrieval as a staged feature reconstruction process, tracking how the reconstruction of a memory trace unfolds in time and space.

## Methods

### Participants

We acquired fMRI data of 37 right-handed, fluent English-speaking participants at the University of Birmingham (26 females, 11 males, mean age (*M_age_*) = 23.33, standard deviation (*SD_age_*) = 3.89, one participant did not indicate their age). The a priori planned sample size for the full EEG-fMRI dataset was *n* = 24 subjects (as in Linde-Domingo et al., 2019). However, due to poor data quality and technical failures in 13 of the simultaneous EEG datasets, additional subjects were recorded leading to a larger sample size for the fMRI data alone. Three subjects were excluded from the fMRI analysis due to failed scanning sequences, and three additional subjects were excluded due to extensive motion within the scanner, exceeding the functional voxel size, such that 31 fMRI datasets remained for analysis. All participants were informed about the experimental procedure, underwent imaging safety screening and signed an informed consent. The research was approved by the STEM ethics committee of the University of Birmingham.

For the fusion analyses, we also included a previously published EEG dataset including the same stimulus set and a nearly identical associative learning and retrieval paradigm (Linde-Domingo et al., 2019). This additional dataset included 24 further participants with a clean, out-of-scanner EEG. For further information on the EEG data sample and the related ethical procedures, we refer to previous work (Linde-Domingo et al., 2019).

### Material

The paradigm was a visual verb-object association task (Linde-Domingo et al., 2019) adapted for fMRI in terms of timing. Stimuli included 128 action verbs and 128 pictures of everyday objects (Linde-Domingo et al., 2019). Action verbs were chosen because they do not elicit concrete object images in themselves but are still easy to associate with the object images. Importantly, all object images existed in two perceptual categories, a coloured photograph from the BOSS database (Brodeur et al., 2010, available on https://sites.google.com/site/bosstimuli/home) or a black line drawing version of a respective photograph created by Linde-Domingo et al. (2019) by means of the free and open source GNU image manipulation software (www.gimp.org). Further each object belonged to one of two conceptual categories, i.e. animate vs inanimate. We selected 128 images per participant according to a fully balanced scheme, such that each combination of perceptual and conceptual categories included the same number of pictures (32 animate-photographs, 32 animate-drawings, 32 inanimate-photographs, 32 inanimate-drawings; Fig. 1a). With respect to the later fusion with the out-of-scanner EEG data, it is important to note that while both experiments used the same set of 128 objects, the same object could appear in different perceptual versions between participants, due to the pseudo-randomised image selection. For example, an image of a camel could be shown as a photograph to one participant, and as a line drawing to another. Action verbs were randomly assigned to images in each participant and were presented together with pictures centrally overlaid on a white background.

### Procedure

Before the session, participants were informed about the experimental procedure and asked to perform a training task block in front of the computer. After completion of the training, the experimental session in the fMRI scanner included four runs with four task blocks each, summing up to a total of 16 task blocks. A typical block in the training and experimental task included the encoding of eight novel associations, a 20 s distractor task, and 16 retrieval trials (two repetitions per association). A 3 min break was included after each fMRI run, in which participants were asked to rest and close their eyes. In total, it took participants approximately 70 min to perform the entire task.

### Encoding

In the encoding phases (Fig. 1b), participants were instructed to study eight novel verb-object pairings in random order. A trial started with a fixation cross presented for a jittered period between 500 and 1500 ms. The action verb was presented for 500 ms before an object was shown for a maximum duration of 5 s. To facilitate learning, participants were instructed to form a vivid visual mental image using the verb-object pairing, and once formed, to press a button with their thumb, which moved the presentation on to the next trial. Importantly, due to the artificial setting where there is no predictive structure to the events, we expected very limited top-down feedback and hence a dominating feedforward sweep during perception early on.

### Distractor

After each encoding phase, participants performed a self-paced distractor task for 20 s, indicating as fast as possible whether each of the consecutively presented numbers on the screen was odd or even, pressing a button with their index or middle finger, respectively. Feedback on the percentage of correct responses was provided at the end of each distractor phase.

### Retrieval

In the retrieval phases, participants were instructed to recall the previously associated objects upon presentation of the corresponding verb (Fig. 1b). Trials started with the presentation of a fixation cross, jittered between 500 and 1500 ms and followed by a previously encoded action verb, presented for 500 ms. Cued with the verb, participants were instructed to recall the paired object within a maximum of 5 s, while a black fixation cross was presented on screen. Note that this maximum duration was shorter than in the original task (Linde-Domingo et al., 2019), which might have caused participants to react faster and possibly explains RT differences between the in-scanner and out-of-scanner datasets (Fig. 1d). Participants were asked to indicate the time point of image recall by button press with their thumb, at which point the black fixation cross turned grey and was presented for an additional 3 s. This retrieval button press was meant to mark the time point of subjective recollection. During the remaining 3 s participants were asked to hold the mental image of the object in mind as vividly as possible. Last, they were asked about the perceptual (Was the object a photograph or a line drawing?) or conceptual (Was the object animate or inanimate?) features of the recalled object, answering with their index or middle finger within a maximum response period of 5 s. During the presentation of the catch question, participants also had the option to indicate with their ring finger that they forgot the features of the corresponding object. Importantly, each encoded stimulus was retrieved twice, once with a perceptual and once with a conceptual question. Trial order was pseudo-random within the first and second set of repetitions, with a minimum of two intervening trials before a specific object was recalled for the second time. The order of catch questions was counterbalanced across repetitions such that half of the associations were first probed with a perceptual question, and the other half was first probed with a conceptual question.

### fMRI data acquisition

FMRI scanning was performed in a 3 Tesla Philips Achieva MRI scanner with a 32-channel SENSE head coil at the Birmingham University Imaging Centre (BUIC, now the Centre for Human Brain Health, CHBH). Slices covered the whole head. T1-weighted anatomical scans were collected with an MP-RAGE sequence (1 mm isotropic voxels, 256 x 256 matrix, 176 slices, no inter-slice gap, repetition time (TR) = 7.5 ms, field of view (FOV) = 256 x 256 x 176 mm, flip angle (FA) = 7°, echo time (TE) = 3.5 ms). T2*-weighted functional images were acquired by a dual-echo EPI pulse sequence with a low specific absorption rate (SAR) to optimize the image quality in regions highly susceptible for distortions by a long readout gradient (3 x 3 x 3.5 mm, 64 x 64 matrix, 34 slices, no inter-slice gap, TR = 2.2 s, full head coverage, FOV = 192 x 192 x 119 mm, FA = 80°, TE1 = 12 ms, TE2 = 34 ms (for dual-echo information see also Halai et al., 2014, 2015; Kirilina et al., 2016). Slices were acquired in a continuous descending fashion. Furthermore, we collected 200 resting state volumes, 50 after each of the four task runs, used for the later combining of dual-echo images with the short and long TEs. During the acquisition of the functional scans, the helium pump was switched off to prevent contamination of the EEG by the compressor artifact at about 20-30 Hz. Stimulus timings were jittered in relation to the acquired scans. The task was presented to participants through a mirror system in the scanner and a JVC SX 21e projector with resolution 1280×1024 at 60 Hz (https://de.jvc.com/). Participantś heads were extra padded to minimize movement artefacts. The stimulus presentation, and the collection of timing and accuracy information, was controlled by scripts written in Matlab 2016a (www.mathworks.com) and Psychophysics Toolbox Version 3 (Brainard, 1997; Kleiner et al., 2007; Pelli, 1997). Responses were logged by NATA response boxes (https://natatech.com/).

### Out-of-scanner EEG data acquisition

As previously stated, a simultaneous EEG dataset was acquired during the fMRI session, but this EEG dataset was too noisy to decode retrieval-related information. We therefore decided to instead use an out-of-scanner EEG dataset for data fusion. This out-of-scanner data, used for all analyses reported here, originated from a previous publication (Linde-Domingo et al., 2019). We give a short description of the EEG data acquisition in the following and of the EEG preprocessing further below. More details can be found in the original publication.

For EEG data collection, 128 electrodes of silver metal and silver chloride were used. Further, an Active-Two amplifier system and the ActiView acquisition software aided data recording (BioSemi, Amsterdam, the Netherlands). Psychophysics Toolbox Version 3 and MATLAB 2014b (www.mathworks.com) were used for task presentation and response collection (Linde-Domingo et al., 2019).

## Analyses

### Behaviour

Behavioural data was analysed by MATLAB R2017b (www.mathworks.com). For the inspection of RTs and accuracies, the behavioural data were preprocessed as follows. For the RT analysis, all trials with incorrect responses to the catch question were removed first. Additionally, catch question RTs of correct trials were removed if they were faster than 200 ms or exceeding the average RT of a participant in a given set of repetitions (first or second cycle) by more than three times the standard deviation (Lifanov et al., 2021; Linde-Domingo et al., 2019). For the analysis of the accuracy data, trials faster than 200 ms and objects with a missing response for either of the two questions were excluded in the corresponding cycle (Lifanov et al., 2021). The same procedures were applied to the data of the out-of-scanner EEG participants, who performed a nearly identical task. However, note that out-of-scanner EEG participants only went through one cycle of retrieval.

### FMRI data preprocessing

We used MATLAB R2017b (www.mathworks.com) and SPM 12 (*Statistical Parametric Mapping*, 2007; http://store.elsevier.com/product.jsp?isbn=9780123725608) for the preprocessing and the analysis of fMRI data. All functional images were first realigned based on three motion and three rotation parameters, unwarped, and slice time corrected to the middle slice in time.

After these initial preprocessing steps, images obtained during the task at two echo times were combined as a weighted average (Poser et al., 2006). Importantly, the relative weights were obtained from the SNR of 200 resting state volumes per echo and its corresponding *TE*

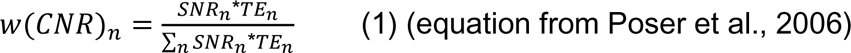

where *w* is the weight for an individual voxel, *SNR_n_*is the signal to noise ratio here calculated as ratio of mean to standard deviation of the given voxel calculated over time at the *n*th echo, and *TE* is the readout time of the *n*th echo.

Weights obtained from the resting state were then used to combine task volumes from both echoes.

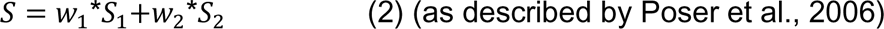

where S is the final signal of an individual voxel over time, calculated by summing the weighted signals of both echoes. These methods and equations were applied as described in work on BOLD contrast optimization by multi-echo sequences (Poser et al., 2006). Combined image structures were written to NIFTI files by Tools for NIfTI and ANALYZE image (https://www.mathworks.com/matlabcentral/fileexchange/8797-tools-for-nifti-and-analyze-image retrieved December 8, 2017, by Shen, 2021).

Anatomical images were segmented, co-registered with the functional images and normalized into a standard MNI template in SPM 12 (*Statistical Parametric Mapping*, 2007; http://store.elsevier.com/product.jsp?isbn=9780123725608). Then, after the combination of functional images, the T2* images were also normalized into MNI space, using the T1-based normalization parameters. Finally, EPI images were smoothed for the univariate GLM analysis with a gaussian spatial filter of 8 mm full width at half maximum (FWHM). Note that multivariate analyses were performed on unsmoothed data in native space (see Weaverdyck et al., 2020).

### ROIs

ROIs were created from templates in MNI space as available in the WFU PickAtlas v3.0 (Lancaster et al., 1997, 2000; Maldjian et al., 2003, 2004; Tzourio-Mazoyer et al., 2002). These ROIs were then fitted to individual brains by applying the reversed T1-based normalization parameters (see previous section) to the ROI masks (as in Chang & Glover, 2010). To define the anatomical masks, we used Brodmann areas (BAs, Brodmann, 1909) from the Talairach Daemon database atlases (Lancaster et al., 1997, 2000) and automated anatomical labelling (AAL, Tzourio-Mazoyer et al., 2002). The anatomical masks used for our analyses included: an early visual mask (V1/V2), consisting of BAs 17 and 18; a human inferior temporal (hIT) mask, consisting of BAs 19 and 37; a temporal pole mask, consisting of superior and middle temporal pole regions as defined by AAL; a medial temporal lobe (MTL) mask, consisting of BAs 28, 34, 35, 36 and AAL rhinal sulcus and parahippocampal gyrus; a superior parietal lobe mask, consisting of BA 7; a lateral (inferior) parietal lobe mask, consisting of BA 39 and 40; and a medial parietal lobe mask, consisting of BA 29 and 30.

### EEG data preprocessing

EEG data (from Linde-Domingo et al., 2019, also available on https://doi.org/10.17605/OSF.IO/327EK) were preprocessed in MATLAB (www.mathworks.com) and the Fieldtrip toolbox version from 3 August, 2017 (Oostenveld et al., 2011); Donders Institute for Brain, Cognition and Behaviour, Radboud University Nijmegen, the Netherlands. See http://www.ru.nl/neuroimaging/fieldtrip). During epoching, different temporal references were used for encoding and retrieval. Encoding epochs were stimulus locked to the onset of the object image, while retrieval timelines were locked to the subjective retrieval button press, in order to observe the reactivation stream leading up to the subjective experience of recollection. These epochs were created with a length of 2 s (−500 before to 1500 ms after object onset) for encoding and 4.5 s (−4 s before to 500 ms after retrieval button press) for retrieval. Line noise was removed by a FIR filter with a band-stop between 48 and 52 Hz. A high-pass filter with a cut-off frequency of 0.1 Hz was applied to remove slow temporal drifts, and a low-pass filter with a cut-off frequency of 100 Hz to remove high-frequency noise. Individual artifactual trials and bad electrodes were rejected manually. Remaining artifacts were removed by independent component analysis (ICA), after which any excluded electrodes were re-introduced by interpolation. The referencing of the data was set to the average across all scalp channels.

After this step, we implemented additional preprocessing steps using MATLAB R2017b (www.mathworks.com) and the Fieldtrip toolbox version from 16 November, 2017 (Oostenveld et al., 2010; Donders Institute for Brain, Cognition and Behaviour, Radboud University Nijmegen, the Netherlands. See http://www.ru.nl/neuroimaging/fieldtrip, version from 16th November, 2017) to prepare data for the specific fusion analyses of this paper. The encoding data was baseline corrected by subtracting the average signal in the pre-stimulus period (−0.2 to −0.1 s) for all post-stimulus time points, separately per electrode. The retrieval data was baseline corrected by whole trial demeaning, since retrieval trials had no obvious, uncontaminated baseline period. The EEG time series data was then downsampled to 128 Hz and temporally smoothed with a moving average with a time window of 40 ms.

### fMRI multivariate analyses

As preparation for the multivariate analyses, we performed a GLM, modelling individual object-specific regressors for encoding, and for each of the two retrieval repetitions separately, adding regressors of no interest for the presentation of verbs, button presses, and catch question onsets, as well as nuisance regressors for head motion, scanner drift and run means (one regressor per variable containing all onsets, see also (Griffiths et al., 2019). We used stick functions locked to the object onset to model the onset of encoding trials and boxcar functions with a duration of 2.5 s locked to the cue onset to model the onset of retrieval trials. The resulting beta weights were transformed into t-values for all subsequent multivariate analyses (Misaki et al., 2010).

### fMRI searchlight LDA

To investigate where in the brain activity patterns differentiated between the two perceptual and the two conceptual categories, we performed a volumetric LDA searchlight analysis on the non-normalized and unsmoothed fMRI data of each participant individually using the searchlight function of the RSA toolbox (Kriegeskorte, 2009; Kriegeskorte et al., 2006, 2008; Kriegeskorte & Kievit, 2013; Nili et al., 2014; https://www.mrc-cbu.cam.ac.uk/methods-and-resources/toolboxes/). The LDA was calculated using the MVPA-Light toolbox (Treder, 2020; https://github.com/treder/MVPA-Light). This was done at each centre voxel of a sphere, where object-specific t-values of the voxels within a 3D searchlight radius of 12 mm were used as feature vectors. Using these feature vectors, we classified perceptual (photo vs drawing) and conceptual (animate vs inanimate) categories by a 5-fold LDA with 5 repetitions, preserving class proportions, separately for encoding and retrieval using all trials. Individual accuracy maps were normalized to MNI space and spatially smoothed with a 10 mm FWHM Gaussian kernel, before second-level t-tests were performed to statistically compare voxel-specific classification accuracies against 50% chance performance. Finally, the results were plotted on an MNI surface template brain.

### fMRI correlation-based RSA

As preparation for the fusion of EEG and fMRI data, we performed a representational similarity analysis (RSA, Kriegeskorte, 2009; Kriegeskorte et al., 2006, 2008; Kriegeskorte & Kievit, 2013; Nili et al., 2014; https://www.mrc-cbu.cam.ac.uk/methods-and-resources/toolboxes/) on the non-normalized and unsmoothed fMRI data. FMRI-based t-maps corresponding to unique objects were arranged in the same order for all participants. As mentioned earlier, object correspondence across participants was only given on the level of object identity (and thus also conceptual category), but not perceptual format. In other words, all participants saw an image of a camel (i.e., an animate object) at some point in the experiment, but the camel could be presented as a photograph to some participants, and as a line drawing to others. For each voxel and its surrounding neighbours within a radius of 12 mm, we extracted object-specific t-value patterns resulting from the appropriate GLM and arranged these as one-dimensional feature vectors. Using these feature vectors, we calculated the Pearson correlation distance (1-*r*) between each pair of objects at each voxel location, separately for encoding and retrieval. The resulting RDM maps were saved and used at a later stage for the searchlight fusion with the EEG data.

In a similar fashion to the searchlight approach, RDMs were also calculated for the pre-defined set of anatomical regions of interest in the non-normalized and unsmoothed individual functional datasets (see section ROIs). The resulting ROI RDMs were used at a later stage for the ROI fusion.

## EEG multivariate analyses

### EEG cross-subjects classifier-based RSA

Our next step was to construct a representational dissimilarity matrix (RDM) for each time point of the EEG recordings, resulting in one timeline representing the similarity structure of our object dataset across all subjects. Cells of the RDM represented the pair-wise discriminability of object pairs, based on a cross-subjects classification of object identity using the MVPA-Light toolbox (Treder, 2020; https://github.com/treder/MVPA-Light). To compute this matrix, we arranged object-specific trials in the same order between all participants, independent of their perceptual format, and up to24 repetitions of each individual object across participants were used for the pairwise classification (Note that after trial removal in the preprocessing, an average of M = 3.09 trials per object were missing in the encoding and M = 3.93 trials per object were missing in the retrieval data). Specifically, we performed a time resolved LDA using as feature vectors the EEG amplitude values from the 128 electrodes, at a given time bin, and participants as repetitions of the same object. To tackle problems of overfitting we used automatic shrinkage regularization as implemented in the MVPA-Light toolbox (Treder, 2020). We used the LDA to classify object identity among each pair of objects at each time bin, again with a 5-fold cross validation, preserving class proportions. The resulting discrimination accuracies were entered in a single time resolved RDM structure representing the dissimilarity between individual objects of our stimulus pool across participants over time. This classification procedure was performed independently for encoding and retrieval.

Before using the EEG-based RDMs for our data fusion, we also wanted to assess statistically how much information about object identity can be decoded from the EEG signals themselves (see Supplements). We therefore first calculated the average classification performance across all pairwise accuracies as a descriptive measure. We then tested if the pairwise accuracies resulting from the ‘real’ classification with correct object labels were significantly larger than the pairwise accuracies that resulted from a classification with permuted object labels. This test was performed in two steps. First, we created 25 classification-based RDMs (same classification procedure as for the ‘real’ RDM time course) but each one with randomly permuted object labels, keeping a given label permutation consistent across time in order to preserve the autocorrelation of the EEG time series. The 25 permutations were averaged to form a single ‘baseline’ RDM time course.

As a second step, we used a cluster-based permutation test to find clusters with temporally extended above-chance decoding accuracy. This cluster-based permutation test compared the t-statistic of each time point resulting from a ‘real matrix’ versus ‘baseline matrix’ t-test, with the t-statistics computed from a ‘real matrix’ versus ‘baseline matrix’ comparison, this time shuffling the ‘real’ and ‘baseline’ cells between the two matrices (again keeping a consistent shuffling across time, independent samples t-test to control for unequal variance, 1000 permutations, cluster-definition threshold of p < .05, as used in previous publications (Cichy et al., 2019; Cichy & Pantazis, 2017; Dobs et al., 2019). Note that the variance in the t-tests for each time bin comes from the pair-wise accuracies contained in the cells of the two (‘real’ and ‘baseline’) classification matrices. All remaining analyses were performed using the (‘real’) classification matrix with the correct object labels.

Next, we tested whether object discriminability systematically differs between objects coming from the same or different conceptual classes (i.e., animate and inanimate objects) within the classification with correct labels. We thus calculated the average accuracies of pairwise classes within and between conceptual categories as a descriptive measure. Using another cluster-based permutation test (paired samples t-test, 1000 permutations, cluster-definition threshold of p < .05), we statistically tested for differences of within-against between-category accuracies over time. This analysis was conducted for conceptual classes only, as there was no correspondence of perceptual class between subjects. The encoding analyses focused on the time period from 0 s s to 1.5 s after object onset, while the retrieval analyses focused on the time period from −3 s to −1 s before retrieval button press. We based this latter time window of interest on previous findings (Linde-Domingo et al., 2019) which indicated perceptual and conceptual decoding peaks prior to −1 s before button press. All cluster permutation tests were implemented by means of the permutest toolbox (https://www.mathworks.com/matlabcentral/fileexchange/71737-permutest retrieved February 16, 2021, by Gerber, 2021; also see Maris & Oostenveld, 2007).

## EEG-fMRI data fusion

### ROI fusion

Two distinct approaches were used for the fusion analyses, with either the fMRI data or the EEG data serving as starting point. The first approach used the fMRI patterns from a given ROI as a starting point, and we thus refer to it as ROI fusion. One RDM was created per participant per ROI, representing the similarity structure (i.e., RDM) in a given anatomical brain region. This ROI RDM was then correlated with the RDM from each time bin of the single, EEG-based RDM time course that represents the pooled similarity structure across subjects (see cross-subject classification of described above and in supplements). Correlations and classification accuracies were Fisher’s z-transformed before the data fusion (as in Griffiths et al., 2019). This analysis resulted in one correlation time course per individual ROI per subject who took part in the fMRI experiment (*n* = 31). The EEG-fMRI correlations were only computed for those cells of the matrix that an individual participant from the fMRI study remembered correctly. To test for statistical significance, we then contrasted the 31 correlation time courses against zero, using a cluster-based permutation test with 1000 permutations and a cluster-definition threshold of p < .05, correcting for multiple comparisons in time as in previous studies (Cichy et al., 2019; Cichy & Pantazis, 2017; Dobs et al., 2019).

To test for a sequential information progression over the ventral visual stream regions, we implemented a linear regression on the cumulative sums of the ROI time courses (based on Michelmann et al., 2019). To do so, we first calculated the cumulative sum of the ROI time courses over the time period from 0 to 1.5 s after image onset for encoding, and from −3 to −1 s before button press for retrieval. For an easier comparison, the cumulative sum of each ROI time course was normalized to an area under the curve that equals 1. For each time point, a linear regression was fitted across the cumulative correlation values of the ROIs within subjects. The resulting slopes of all subjects were then tested against zero in a one-sided cluster-based permutation test (with 1000 permutations and a cluster-definition threshold of p < .05). This method enabled us to test for a forward stream during encoding and a backward stream during retrieval (Fig. 4e-f). An example of the rationale is illustrated in figure 4. In the case of a forward stream, the ROIs along the ventral visual stream activate sequentially from early towards late regions. Therefore, earlier regions (e.g. V1-hIT) show a higher cumulative sum than later regions (e.g. temporal pole - MTL) at 0.5 s after stimulus onset (and other time points). A linear fit across ROIs at 0.5 s will therefore show a significant negative slope. According to this rationale, a backward stream across the same regions would result in a significant positive slope. Since we expected a forward stream at encoding and a backward stream at retrieval, we used one-sided tests to confirm if the slope differs from zero (< 0 at encoding, > 0 at retrieval). The sequential ordering of our ROIs was based on the extensive literature describing the hierarchical organisation of the ventral visual stream in anatomical and functional neuroimaging studies (e.g. Cichy et al., 2016, 2017; Felleman & Essen, 1991).

### Searchlight fusion

The second, complementary fusion approach offers higher spatial resolution and a whole-brain perspective on how encoding and retrieval patterns emerge over time. Here, we used the EEG-based RDMs from each time point as starting point, and then searched for matching similarity structures across the entire brain, using a volumetric searchlight analysis on each individual’s fMRI data (Kriegeskorte, 2009; Kriegeskorte et al., 2006, 2008; Kriegeskorte & Kievit, 2013; Nili et al., 2014). We refer to this method as a (time-resolved) searchlight fusion. A second-order correlation was computed between the (classification-based) EEG RDM from each time bin and the (correlation-based) fMRI RDMs for each centre voxel and its neighbours within a searchlight radius of 3 voxels, separately for encoding and retrieval. This analysis results in a “fusion movie” (i.e., a time-resolved brain map) for each participant who took part in the fMRI experiment. The searchlight fusion for retrieval data was performed for correct trials within the fMRI data only (same as above). The fused data was Fisher’s z-transformed. The searchlight fusion was performed using the Python Representational Similarity Analysis (rsatoolbox) toolbox (https://rsatoolbox.readthedocs.io/en/latest/, https://github.com/rsagroup/rsatoolbox) which works on Python (van Rossum, 1995), using the sys and os module, SciPy (Virtanen et al., 2020), NumPy (Harris et al., 2020), and nibabel (https://nipy.org/nibabel/; https://github.com/nipy/nibabel/releases).

To test for significant EEG-fMRI pattern similarity at individual voxels, we normalized the correlation maps to MNI space, smoothed them with a 10 mm FWHM Gaussian kernel and then tested them in a one-sample t-test against zero at each single time bin. The t-test included a spatial maximal permuted statistic correction combined with a threshold free cluster enhancement (Nichols & Holmes, 2002; Smith & Nichols, 2009). This cluster-based analysis was performed using the toolbox MatlabTFCE (http://markallenthornton.com/blog/matlab-tfce/) with 1000 permutations, a height exponent of 2, an extent exponent of 0.5, a connectivity parameter of 26 and a step number for cluster formation of .1 as suggested by Smith & Nichols (2009). The analysis resulted in time-resolved spatial t-maps, depicting clusters of significant EEG-fMRI correlations, created by Tools for NIfTI and ANALYZE image (https://www.mathworks.com/matlabcentral/fileexchange/8797-tools-for-nifti-and-analyze-image retrieved December 8, 2017, by Shen, 2021).

### Figures

Figures were created using SPM 12 (*Statistical Parametric Mapping*, 2007; http://store.elsevier.com/product.jsp?isbn=9780123725608), the RainCloud plots Version 1.1 (https://github.com/RainCloudPlots/RainCloudPlots, Allen et al., 2019; Whitaker et al., 2019), Inkscape 1.0.1 (https://inkscape.org/), WFU PickAtlas v3.0 (Maldjian et al., 2003, 2004), MRIcron (https://www.nitrc.org/projects/mricron, www.mricro.com, Rorden & Brett, 2000) and a colin 27 average brain template (http://www.bic.mni.mcgill.ca/ServicesAtlases/Colin27, Holmes et al., 1998).

## Analyses that did not work

### 1) Data fusion with the in-scanner EEG

The initial plan for this project was to fuse the fMRI data with the EEG data acquired during scanning. Using the simultaneous EEG data for fusion, the spatio-temporal mapping could then have been performed on a trial-by-trial basis within participants. However, neither a time-resolved RSA nor a perceptual or conceptual LDA classification on the EEG data showed interpretable results, likely due to extensive noise in the in-scanner EEG data.

### 2) Data fusion with correlation-based EEG RDMs

Having decided to use the out-of-scanner EEG dataset for the data fusion, we first attempted to perform the fMRI-EEG fusion using correlation-based RDMs calculated from the EEG data (see Cichy & Oliva, 2020). To compute these RDMs, we arranged object-specific trials in the same order between all participants, independent of their perceptual format, identical to the arrangement in the fMRI data. We then performed a time-resolved RSA using EEG amplitude values from the 128 electrodes, at a given time bin, as feature vectors. Specifically, we calculated the pairwise distance (1-Pearson correlation (*r*)) between each pair of objects. However, inspecting the average overall similarity and contrasting the average similarity within versus between conceptual classes did not show interpretable results. This is most likely due to too unreliable and time-varying single-trial estimates from the EEG data. Following this initial analysis, we thus used the cross-subject, classification-based RDMs that produced more stable results on the EEG data itself.

## Data availability

The EEG data (Linde-Domingo et al., 2019) can be found under https://doi.org/10.17605/OSF.IO/327EK. fMRI data can only partly be made publicly available to guarantee full anonymity in accordance with participants’ consent. Group-level data and non-identifiable individual-level data can be found on the Open Science Framework with the identifier https://doi.org/10.17605/OSF.IO/2T7SN. Consent for the reuse of data does not include commercial research.

## Code availability

The custom code used in this study is available on the Open Science Framework with the identifier https://doi.org/10.17605/OSF.IO/2T7SN. Additional code associated with the EEG data (Linde-Domingo et al., 2019) can be found under https://doi.org/10.17605/OSF.IO/327EK.

## Acknowledgements and Funding

We thank, Simrandeep Cheema, Dagmar Fraser, and Nina Salman for help with the data collection. Further we thank Karen Mullinger for useful analytical advice. This work was supported by a European Research Council (ERC) Starting Grant StG-2016-715714 awarded to Maria Wimber, and a scholarship from the Midlands Integrative Biosciences Training Partnership (MIBTP) awarded to Juan Linde-Domingo. The funders had no role in study design, data collection and interpretation, or the decision to submit the work for publication.

## Author contributions

J.L., J.L.D. and M. Wimber designed the experiments. J.L. and J.L.D. conducted the experiments. B.G. and C.F. helped with the EEG-fMRI acquisition. J.L. and J.L.D preprocessed the data. J.L. analysed the data. All authors contributed to the analyses. J.L. and M. Wimber wrote the manuscript. All authors contributed to reviewing and editing.

## Ethics

Human subjects: All participants gave informed written consent. The research (ethics code ERN_11-0429AP70) was approved by the STEM ethics committee of the University of Birmingham.

## Supplementary file 1

### fMRI searchlight LDA

**Table 3.**
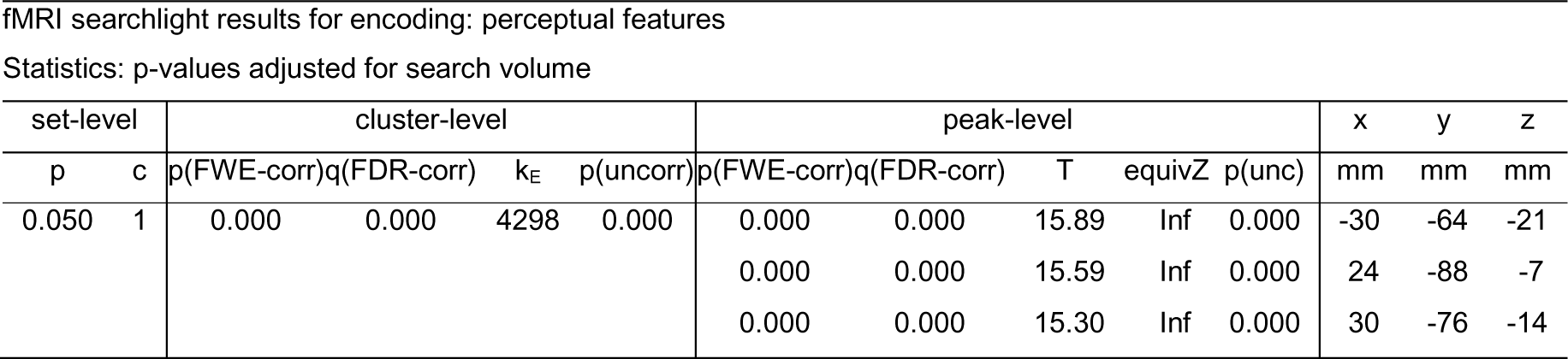
fMRI searchlight results for encoding: perceptual features.

| set-level |  | cluster-level |  |  |  | peak-level |  |  |  |  | x | y | z |
| --- | --- | --- | --- | --- | --- | --- | --- | --- | --- | --- | --- | --- | --- |
| p | c | p(FWE-corr) | q(FDR-corr) | k<sub>E</sub> | p(uncorr) | p(FWE-corr) | q(FDR-corr) | T | equivZ | p(unc) | mm | mm | mm |
| 0.050 | 1 | 0.000 | 0.000 | 4298 | 0.000 | 0.000 | 0.000 | 15.89 | Inf | 0.000 | -30 | -64 | -21 |
|  |  |  |  |  |  | 0.000 | 0.000 | 15.59 | Inf | 0.000 | 24 | -88 | -7 |
|  |  |  |  |  |  | 0.000 | 0.000 | 15.30 | Inf | 0.000 | 30 | -76 | -14 |

Statistics: p-values adjusted for search volume

**Table 4.** fMRI searchlight results for encoding: conceptual features.

| set-level |  | cluster-level |  |  |  | peak-level |  |  |  |  | x | y | z |
| --- | --- | --- | --- | --- | --- | --- | --- | --- | --- | --- | --- | --- | --- |
| p | c | p(FWE-corr) | q(FDR-corr) | k<sub>E</sub> | p(uncorr) | p(FWE-corr) | q(FDR-corr) | T | equivZ | p(unc) | mm | mm | mm |
| 0.000 | 3 | 0.000 | 0.000 | 7638 | 0.000 | 0.000 | 0.000 | 20.98 | Inf | 0.000 | 45 | -73 | 4 |
|  |  |  |  |  |  | 0.000 | 0.000 | 16.79 | Inf | 0.000 | -39 | -79 | 7 |
|  |  |  |  |  |  | 0.000 | 0.000 | 16.71 | Inf | 0.000 | 42 | -61 | -21 |
|  |  | 0.000 | 0.007 | 212 | 0.005 | 0.003 | 0.054 | 6.12 | 4.89 | 0.000 | 54 | 26 | 14 |
|  |  | 0.033 | 0.645 | 5 | 0.645 | 0.019 | 0.383 | 5.24 | 4.38 | 0.000 | -6 | 68 | 21 |

**Table 5.**
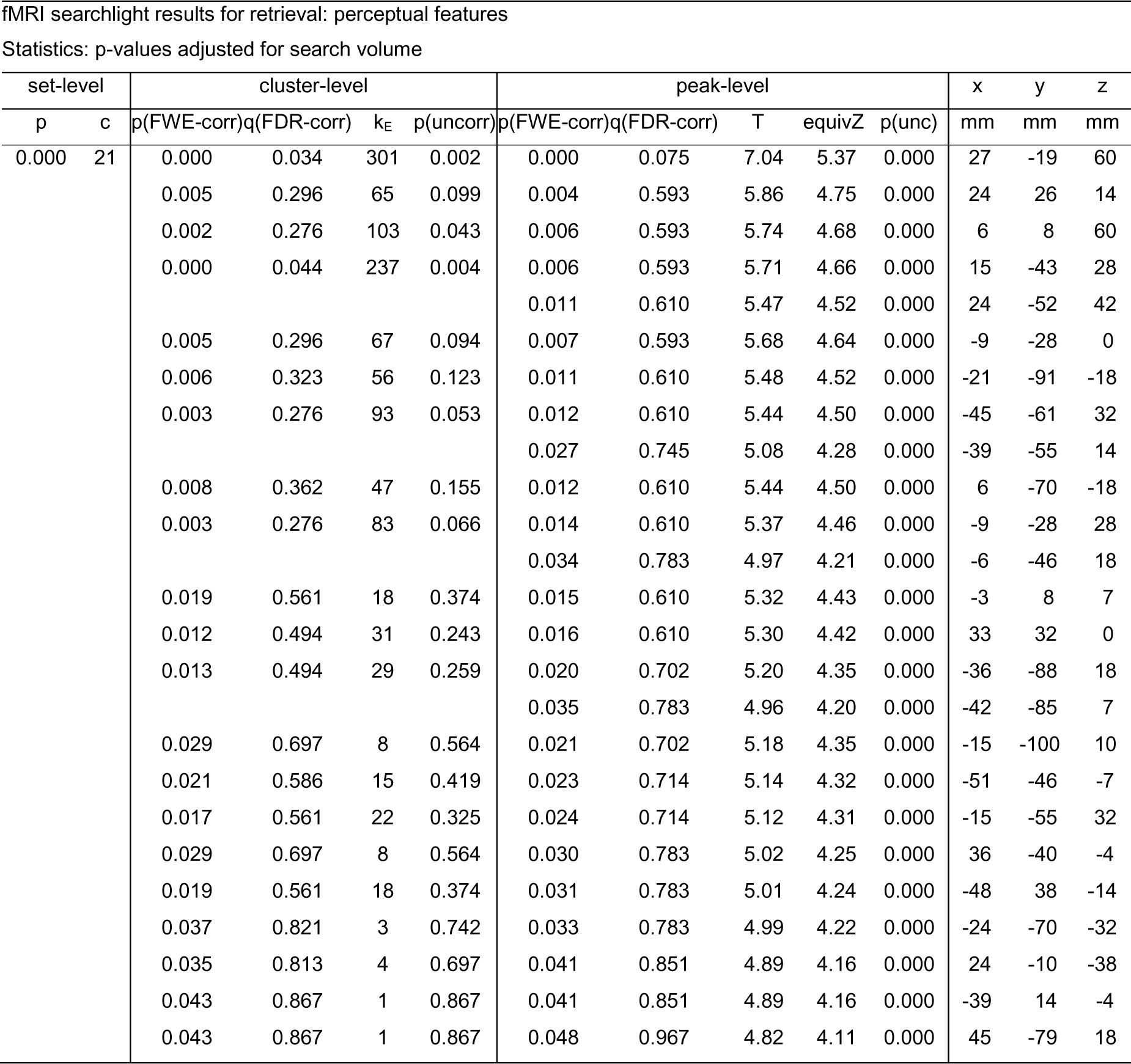
fMRI searchlight results for retrieval: perceptual features.

| set-level |  | cluster-level |  |  |  | peak-level |  |  |  |  | x | y | z |
| --- | --- | --- | --- | --- | --- | --- | --- | --- | --- | --- | --- | --- | --- |
| p | c | p(FWE-corr) | q(FDR-corr) | k<sub>E</sub> | p(uncorr) | p(FWE-corr) | q(FDR-corr) | T | equivZ | p(unc) | mm | mm | mm |
| 0.000 | 21 | 0.000 | 0.034 | 301 | 0.002 | 0.000 | 0.075 | 7.04 | 5.37 | 0.000 | 27 | -19 | 60 |
|  |  | 0.005 | 0.296 | 65 | 0.099 | 0.004 | 0.593 | 5.86 | 4.75 | 0.000 | 24 | 26 | 14 |
|  |  | 0.002 | 0.276 | 103 | 0.043 | 0.006 | 0.593 | 5.74 | 4.68 | 0.000 | 6 | 8 | 60 |
|  |  | 0.000 | 0.044 | 237 | 0.004 | 0.006 | 0.593 | 5.71 | 4.66 | 0.000 | 15 | -43 | 28 |
|  |  |  |  |  |  | 0.011 | 0.610 | 5.47 | 4.52 | 0.000 | 24 | -52 | 42 |
|  |  | 0.005 | 0.296 | 67 | 0.094 | 0.007 | 0.593 | 5.68 | 4.64 | 0.000 | -9 | -28 | 0 |
|  |  | 0.006 | 0.323 | 56 | 0.123 | 0.011 | 0.610 | 5.48 | 4.52 | 0.000 | -21 | -91 | -18 |
|  |  | 0.003 | 0.276 | 93 | 0.053 | 0.012 | 0.610 | 5.44 | 4.50 | 0.000 | -45 | -61 | 32 |
|  |  |  |  |  |  | 0.027 | 0.745 | 5.08 | 4.28 | 0.000 | -39 | -55 | 14 |
|  |  | 0.008 | 0.362 | 47 | 0.155 | 0.012 | 0.610 | 5.44 | 4.50 | 0.000 | 6 | -70 | -18 |
|  |  | 0.003 | 0.276 | 83 | 0.066 | 0.014 | 0.610 | 5.37 | 4.46 | 0.000 | -9 | -28 | 28 |
|  |  |  |  |  |  | 0.034 | 0.783 | 4.97 | 4.21 | 0.000 | -6 | -46 | 18 |
|  |  | 0.019 | 0.561 | 18 | 0.374 | 0.015 | 0.610 | 5.32 | 4.43 | 0.000 | -3 | 8 | 7 |
|  |  | 0.012 | 0.494 | 31 | 0.243 | 0.016 | 0.610 | 5.30 | 4.42 | 0.000 | 33 | 32 | 0 |
|  |  | 0.013 | 0.494 | 29 | 0.259 | 0.020 | 0.702 | 5.20 | 4.35 | 0.000 | -36 | -88 | 18 |
|  |  |  |  |  |  | 0.035 | 0.783 | 4.96 | 4.20 | 0.000 | -42 | -85 | 7 |
|  |  | 0.029 | 0.697 | 8 | 0.564 | 0.021 | 0.702 | 5.18 | 4.35 | 0.000 | -15 | -100 | 10 |
|  |  | 0.021 | 0.586 | 15 | 0.419 | 0.023 | 0.714 | 5.14 | 4.32 | 0.000 | -51 | -46 | -7 |
|  |  | 0.017 | 0.561 | 22 | 0.325 | 0.024 | 0.714 | 5.12 | 4.31 | 0.000 | -15 | -55 | 32 |
|  |  | 0.029 | 0.697 | 8 | 0.564 | 0.030 | 0.783 | 5.02 | 4.25 | 0.000 | 36 | -40 | -4 |
|  |  | 0.019 | 0.561 | 18 | 0.374 | 0.031 | 0.783 | 5.01 | 4.24 | 0.000 | -48 | 38 | -14 |
|  |  | 0.037 | 0.821 | 3 | 0.742 | 0.033 | 0.783 | 4.99 | 4.22 | 0.000 | -24 | -70 | -32 |
|  |  | 0.035 | 0.813 | 4 | 0.697 | 0.041 | 0.851 | 4.89 | 4.16 | 0.000 | 24 | -10 | -38 |
|  |  | 0.043 | 0.867 | 1 | 0.867 | 0.041 | 0.851 | 4.89 | 4.16 | 0.000 | -39 | 14 | -4 |
|  |  | 0.043 | 0.867 | 1 | 0.867 | 0.048 | 0.967 | 4.82 | 4.11 | 0.000 | 45 | -79 | 18 |

**Table 6.**
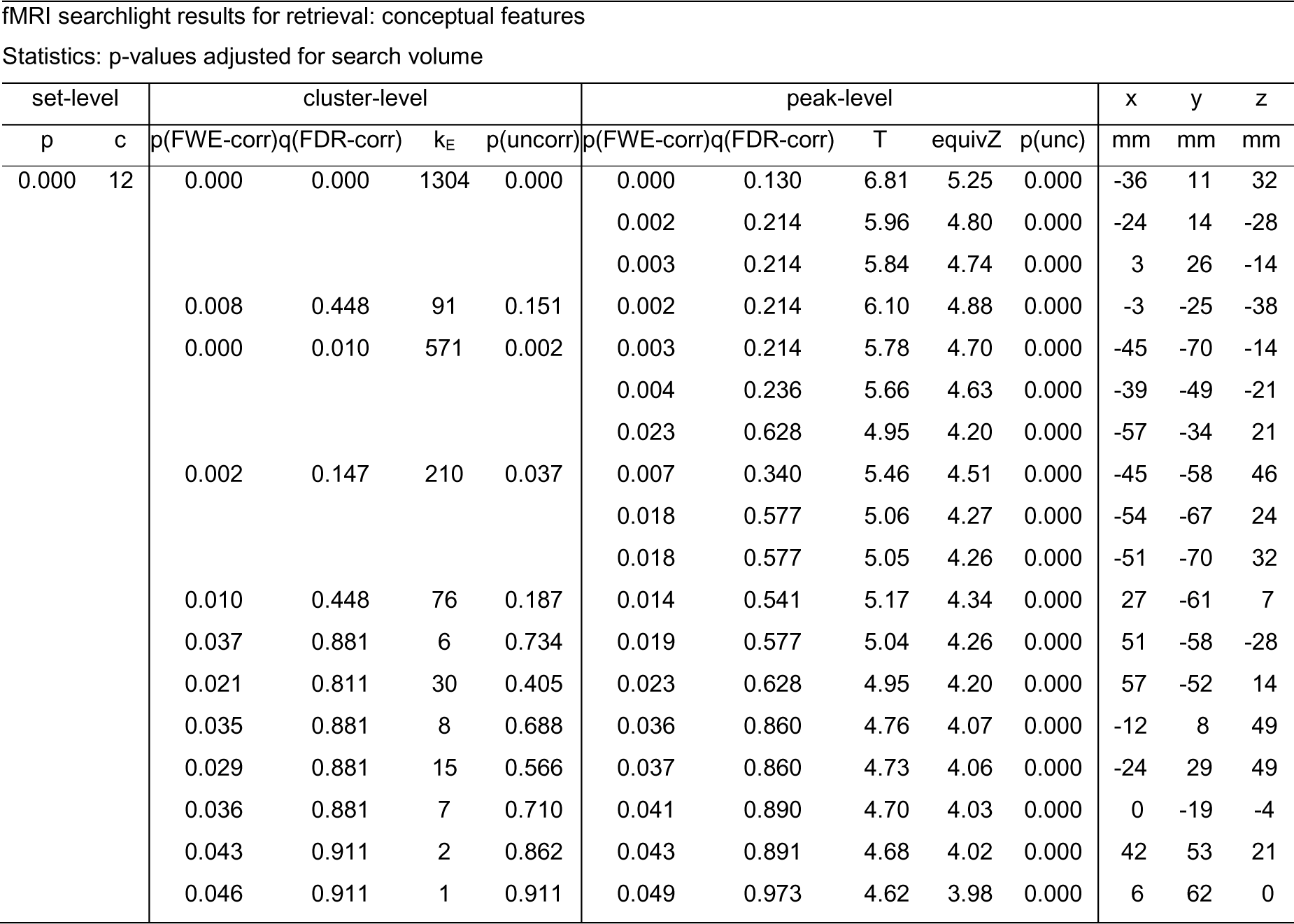
fMRI searchlight results for retrieval: conceptual features.

## Supplementary file 2

### fMRI univariate analyses

We first performed a univariate GLM analysis and subsequent t-contrasts on the fMRI data to reveal activation differences between the two perceptual and the two conceptual categories (Fig. 2, Suppl. Fig. 1). The GLM included the four main regressors (drawing-animate, photograph-animate, drawing-inanimate, and photograph-inanimate), with separate regressors for encoding, first retrieval, and second retrieval, using all trials. We used stick functions locked to the object onset to model the onset of encoding trials and boxcar functions with a duration of 2.5 s locked to the cue onset to model the onset of retrieval trials. We added one regressor each for the presentation of verbs, button presses, and perceptual and conceptual catch questions as well as nuisance regressors for head motion, scanner drift and run means. After computing the GLM for each subject, a sample-level ANOVA was computed with the perceptual (photograph versus line drawing) and conceptual (animate versus inanimate) dimensions as within-subjects factors. Planned comparisons contrasting photographs versus drawings and animate versus inanimate objects were carried out in subsequent t-contrasts separately at encoding and retrieval for all trials. The t-contrasts for retrieval trials were performed using both the first and second retrieval trials together.

### fMRI univariate results

At encoding, photographs and line-drawings showed average activity differences primarily in ventral visual regions (see Fig. 2, Suppl. Fig. 1). Photographs elicited significantly higher BOLD responses in early visual regions, such as V1, V2 and fusiform gyrus, (*t*(30) = 4.56, *p* < .05 (FWE)). When contrasting conceptual categories, animate objects were associated with significantly higher activity in V1, V2, but also fusiform gyrus, inferior temporal gyrus and visuomotor cortex (*t*(30) = 4.56; *p* < .05 (FWE)), while inanimate objects showed a stronger signal in middle occipital gyrus and fusiform gyrus (*t*(30) = 4.56; *p* < .05 (FWE)). These contrasts thus show the expected tendency that perceptual categories differed in regions earlier along the ventral visual stream than conceptual categories, in line with the multivariate results reported in the main results.

During retrieval, no cortical voxels survived family-wise error correction (p < .05 FWE) when contrasting perceptual or conceptual categories.

**Figure 2 - Supplemental Figure 1.**
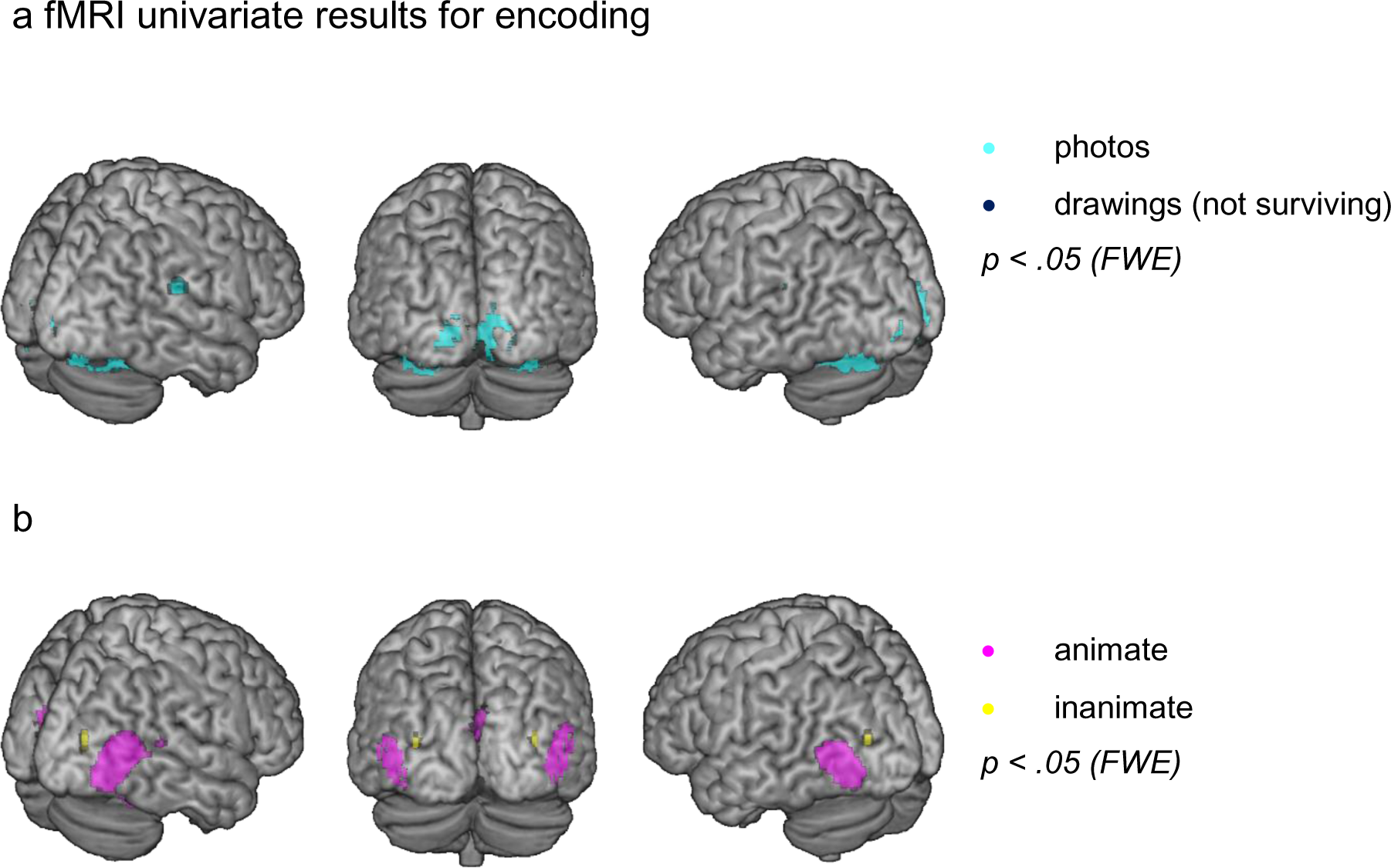
Univariate results. Second-level t-contrasts for encoding. a) Cyan: Photograph > drawing, green: drawing > photograph. b) Magenta: animate > inanimate, yellow: animate< inanimate. All contrasts are thresholded at t(30) = 4.56, p < .05 (FWE). N = 31 independent subjects. Figure made using MRIcron (https://www.nitrc.org/projects/mricron, www.mricro.com, Rorden & Brett, 2000) and a colin 27 average brain template (http://www.bic.mni.mcgill.ca/ServicesAtlases/Colin27, Holmes et al., 1998; Copyright (C) 1993–2009 Louis Collins, McConnell Brain Imaging Centre, Montreal Neurological Institute, McGill University).

**Table 7.**
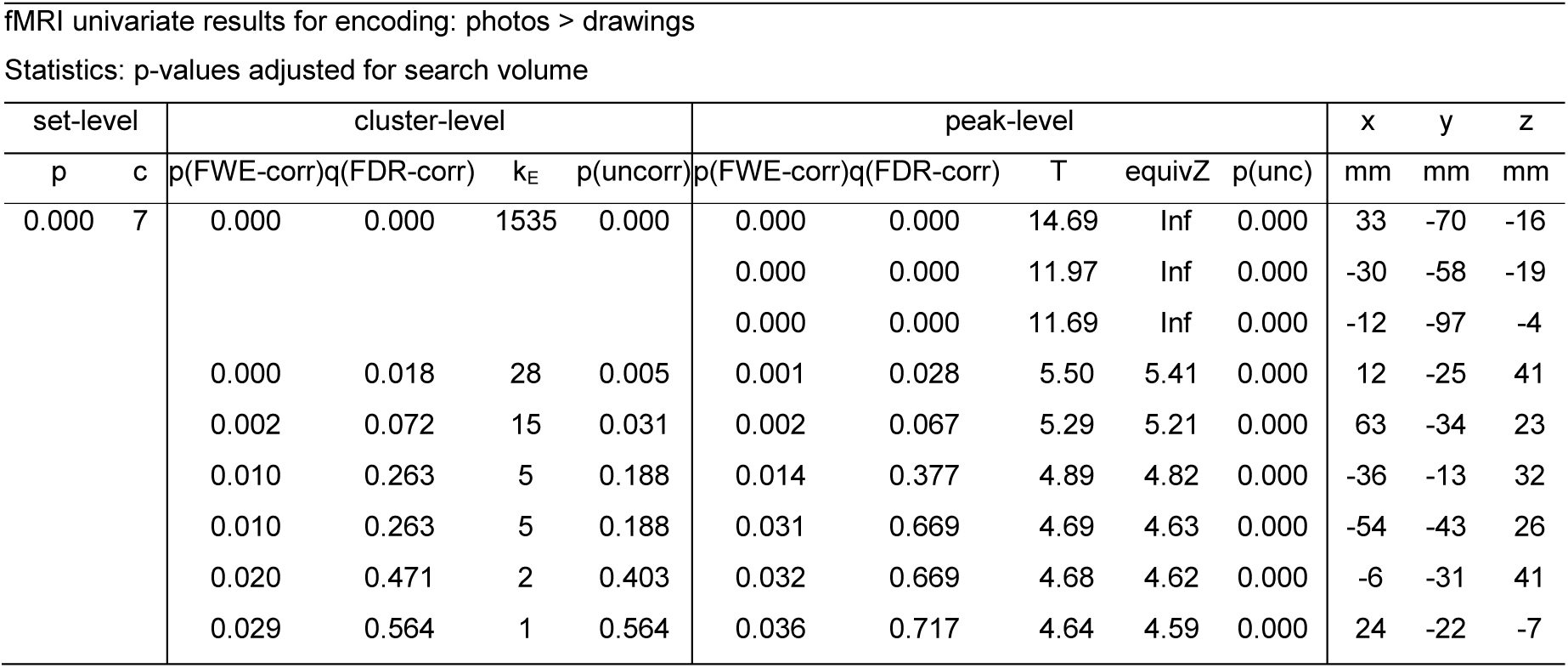
fMRI univariate results for encoding: photos > drawings.

**Table 8.** fMRI univariate results for encoding: animate > inanimate.

| set-level |  | cluster-level |  |  |  | peak-level |  |  |  |  | x | y | z |
| --- | --- | --- | --- | --- | --- | --- | --- | --- | --- | --- | --- | --- | --- |
| p | c | p(FWE-corr) | q(FDR-corr) | k<sub>E</sub> | p(uncorr) | p(FWE-corr) | q(FDR-corr) | T | equivZ | p(unc) | mm | mm | mm |
| 0.000 | 5 | 0.000 | 0.000 | 409 | 0.000 | 0.000 | 0.000 | 14.25 | Inf | 0.000 | 51 | -70 | 5 |
|  |  |  |  |  |  | 0.000 | 0.000 | 10.17 | Inf | 0.000 | 42 | -49 | -22 |
|  |  |  |  |  |  | 0.001 | 0.040 | 5.40 | 5.31 | 0.000 | 54 | -43 | 14 |
|  |  | 0.000 | 0.000 | 170 | 0.000 | 0.000 | 0.000 | 11.59 | Inf | 0.000 | -48 | -76 | 5 |
|  |  | 0.004 | 0.105 | 9 | 0.084 | 0.000 | 0.000 | 6.73 | 6.56 | 0.000 | -39 | -46 | -22 |
|  |  | 0.000 | 0.000 | 145 | 0.000 | 0.000 | 0.003 | 5.97 | 5.86 | 0.000 | 9 | -76 | 8 |
|  |  |  |  |  |  | 0.003 | 0.083 | 5.22 | 5.14 | 0.000 | -6 | -88 | 20 |
|  |  |  |  |  |  | 0.009 | 0.204 | 4.99 | 4.92 | 0.000 | 6 | -79 | 29 |
|  |  | 0.006 | 0.123 | 7 | 0.123 | 0.010 | 0.213 | 4.96 | 4.89 | 0.000 | 9 | -52 | 50 |

Statistics: p-values adjusted for search volume

**Table 9.**
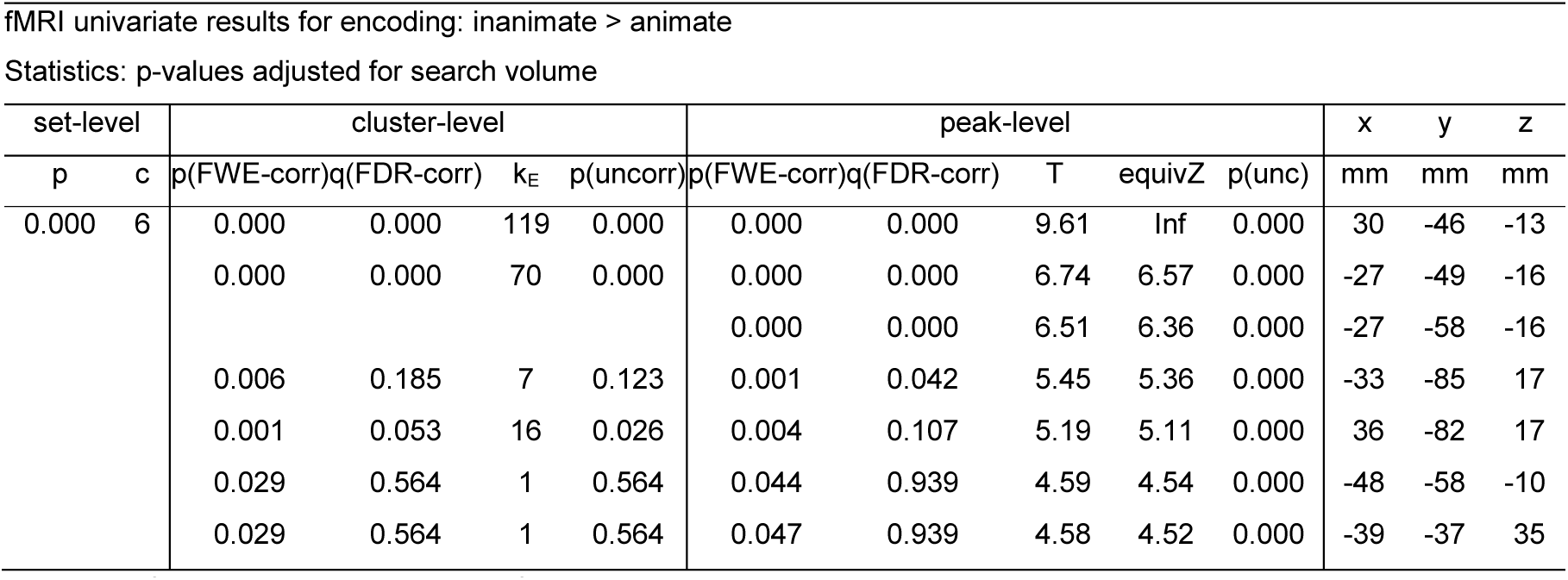
fMRI univariate results for encoding: inanimate > animate.

## Supplementary file 3

### EEG multivariate results

#### EEG cross-subjects classifier-based RSA

Plots in Fig. 4, suppl. Fig. 1 show the time-resolved decoding accuracy from the cross-subjects classifier, once in grey averaged across all object pairings in the RDM, and in magenta split into the pairings belonging to the same (within) or different (between) conceptual classes, all with corresponding statistics (see methods).

During encoding, individual object decoding accuracy started increasing gradually from approximately 25 ms after stimulus onset, with a first smaller peak around 127 ms (p < .05 (cluster)), and a second temporally extended cluster from 236 ms until the end of the trial (p < .05 (cluster)). Moreover, objects from conceptually different classes (animate vs inanimate) were classified significantly better than objects from the same class, starting from 220 ms after stimulus onset until 1.3 s (p < .05 (cluster)). Given the object recognition literature (Cichy et al., 2014), it was surprising that the accuracy of object-identity decoding was comparatively low (though still significant) within the first 100 ms compared to the time period after 200 ms. This might reflect the fact that across participants, the objects corresponded in terms of image content but not perceptual format, potentially suppressing some low-level similarities and thus making classification on early visual features more difficult. Having said that, the photographic and drawn object versions overlapped in shape, orientation and disparity, preserving at least some low-level features, and a classifier should be able to use the neural signals corresponding to these features to discriminate individual objects. In fact, the searchlight fusion shows major correlation clusters during encoding in early visual areas (Fig. 5), indicating that some low-level features contribute to decoding performance at these early time points.

During retrieval, overall cross-subject classification accuracies remained relatively low in comparison to encoding, showing a peak decoding accuracy of 51 % approximately −2.8 s before the retrieval button press. Five significant accuracy clusters were found with peaks located at times −2.74 s, −2.63 s, −2.56 s, −2.16 s and −1.47 s relative to the time of subjective recollection (all p < .05 (cluster)). When comparing classification accuracy for conceptually different, between-class object pairs to within-class pairs, at no time point did this comparison survive cluster correction. The maximum between-versus within-class difference was identified about −2.5 s before the retrieval button press, though at a very liberal threshold (p < .01 (uncorr.)).

**Figure 4 - Supplemental Figure 1.**
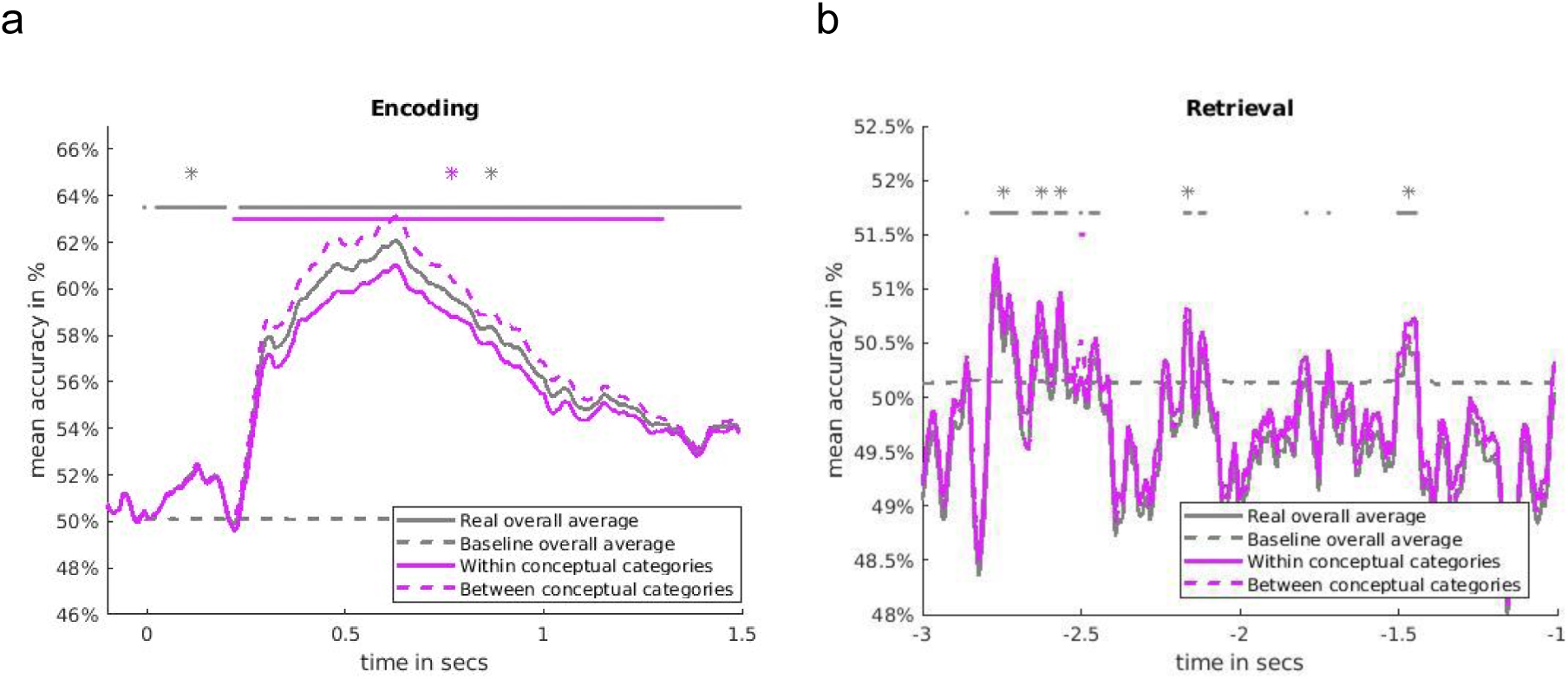
Average accuracy of EEG-based classification of object identity over time (grey), and average classification accuracy within-versus between conceptual classes (magenta) during a) encoding and b) retrieval. At encoding, time point 0 s marks the object onset. At retrieval, time point 0 s marks the button press, but is not included in figure as it does not lie within the time window of interest (see methods). Solid grey line represents average accuracy across the entire RDM, dashed grey line represents baseline average, and grey markers indicate significant overall classification accuracy against chance through random label permutation, compared by independent-sample t-tests. Solid magenta line represents the average accuracy within conceptual categories, dashed magenta line represents average accuracy between conceptual categories, and magenta markers indicate significant between-versus within conceptual classes with correct labels, compared by paired-sample t-tests. Significant decoding accuracy is indicated by points (p < .01 uncorrected) and asterisk (p < .05 cluster). See Methods for details on cluster correction. N = 24 independent subjects.

