## Supplementary figures and images for "Reconstructing Spatio-Temporal Trajectories of Visual Object Memories in the Human Brain"

### Supplemental Figure 2-1

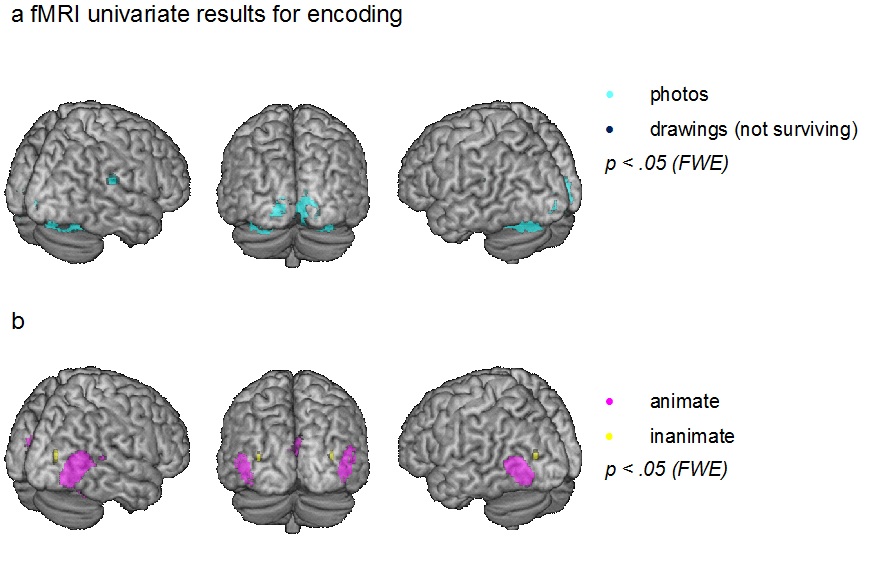

### Supplemental Figure 4-1

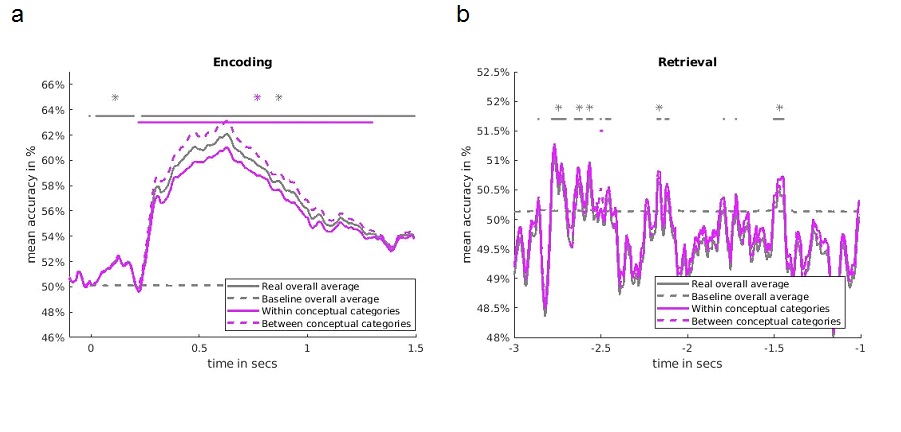
